# Gene genealogies in diploid populations evolving according to sweepstakes reproduction

**DOI:** 10.64898/2026.01.15.699673

**Authors:** Bjarki Eldon

## Abstract

Recruitment dynamics, or the distribution of the number of offspring among individuals, is central for understanding ecology and evolution. Sweepstakes reproduction (when the offspring number distribution has a heavy right-tail) may characterize the recruitment dynamics of highly fecund natural populations. Sweepstakes reproduction can induce jumps in type frequencies, and multiple mergers in gene genealogies of sampled gene copies. Here, we consider gene genealogies in diploid panmictic populations evolving in a random environment. The heavy-tailed offspring number distribution is generated by mechanisms not involving natural selection, such as in chance matching of broadcast spawning with favourable environmental conditions. Our model of sweepstakes reproduction extends the one considered by Schweinsberg [2003] by applying an upper bound to the number of potential offspring of any given parent pair. Depending on the stated bound, the gene genealogies are in the domain of attraction of the Kingman coalescent, or specific familes of continuous-time Beta- or Poisson-Dirichlet simultaneous multiple-merger coalescents. The gene genealogies in a large population are viewed on a timescale proportional to at least *N/* log *N* generations; *N* is proportional to the population size (when constant). Incorporating deterministic population size changes leads to time-changed coalescents; the time-change is independent of the skewness of the offspring-number distribution. Using simulations, we show that gene genealogies in finite populations are not well approximated by the coalescent trees. Simulation results also indicate that quenched (conditioned on the population ancestry) and annealed gene genealogies in finite populations are not in good agreement whenever the skewness of the offspring number distribution is increased.

**Significance Statement:** It has been suggested that highly fecund natural populations characterised by Type III survivorship may evolve according to sweepstakes reproduction, meaning high skewness in the offspring number distribution. Sweepstakes reproduction may affect the evolution of the population in a number of ways. Here, we consider gene genealogies in diploid panmictic populations with a heavy-tailed offspring number distribution, and evolving in a random environment. We derive simultaneous multiple-merger coalescents, time-changed when the population size varies in time. We use simulations to study patterns of genetic variation as predicted by the resulting coalescents, and to compare quenched (conditioning on the ancestral relations of the gene copies in the population) and annealed gene genealogies. Our results are relevant for inferring sweepstakes.

## 1 Introduction

Inferring evolutionary histories of natural populations is one of the main aims of population genetics. Inheritance, or the transfer of gene copies from a parent to an offspring, is the characteristic of organisms that makes inference possible. Inheritance leaves a ‘trail’ of ancestral relations. The structure of the trail shapes the pattern of genetic variation observed in a sample. By modeling the random (unknown) ancestral relations of sampled gene copies, one hopes to be able to distinguish between evolutionary histories by comparing model predictions to population genetic data. This sample-based approach to inference forms a framework for deriving efficient inference methods [Donnelly and Tavaré, 1995, Bertoin, 2006, Wakeley, 2009, Berestycki, 2009].

Recruitment dynamics, or individual recruitment success (the offspring number distribution) is central to ecology and evolution [Eldon, 2020, Hedgecock and Pudovkin, 2011]. In the absence of natural selection and complex demography the offspring number distribution is a deciding factor in how the sample trees look like, and therefore for predictions about data. Models such as the Wright-Fisher model [Wright, 1931, Fisher, 1923], in which large families (with numbers of offspring proportional to the population size) occur only with negligible probability in an arbitrarily large population, are commonly used as offspring number distributions. However, such ‘small family’ models may be a poor choice for highly fecund populations [Hedgecock and Pudovkin, 2011, Árnason et al., 2023].

Highly fecund natural populations are diverse and widely found [Eldon, 2020]. By ‘high fecundity’ we mean that individuals have the capacity to produce numbers (at least) proportional to the population size of ‘potential’ offspring (the surviving offspring are sampled from the pool of potential offspring). It has been suggested that the evolution of highly fecund populations may be characterised by high variance in the offspring number distribution [sweepstakes reproduction; Avise et al., 1988, Palumbi and Wilson, 1990, Li and Hedgecock, 1998, Hedgecock et al., 1982, Hedgecock, 1994, Beckenbach, 1994, Árnason, 2004, Árnason and Halldórsdóttir, 2015, Vendrami et al., 2021, Árnason et al., 2023]. We will use the term ‘sweepstakes reproduction’ for when there occasionally (randomly occurring, and without involving natural selection) is an increased chance of large families with numbers of surviving offspring proportional to the population size.

The evolution of populations evolving according to sweepstakes reproduction may be different from the evolution of populations not characterised by sweepstakes. Forward-in-time processes (in the form of Fleming-Viot measure-valued diffusions [Fleming and Viot, 1979, Ethier and Kurtz, 1993]) describing the evolution of type frequencies in populations characterised by sweepstakes admit jumps [Birkner and Blath, 2009]. Gene genealogies describing the random ancestral relations of gene copies sampled from populations evolving according to sweepstakes are characterised by multiple mergers, where a random number of ancestral lineages merges in one or more groups whenever mergers occur [Donnelly and Kurtz, 1999, Pitman, 1999, Sagitov, 1999, Schweinsberg, 2000, Sagitov, 2003, Möhle and Sagitov, 2001, Möhle and Sagitov, 2003, Huillet and Möhle, 2011, 2013]. Strong positive selection inducing recurrent selective sweeps, recurrent strong bottlenecks, and recurrent extinction and recolonisation in subdivided populations, are additional examples of mechanisms generating multiple-mergers in gene genealogies [Durrett and Schweinsberg, 2005, Birkner et al., 2009, Eldon and Wakeley, 2006, Taylor and Véber, 2009]. Loosely speaking, sweepstakes introduce jumps to the evolution of the population, where ‘jumps’ refer to multiple mergers of ancestral lineages, and discontinuous changes in type frequencies. We are only beginning to understand *(i)* if one can distinguish between the mechanisms producing jumps using population genetic data [Birkner et al., 2009, Hoban et al., 2013, Freund et al., 2023, Johri et al., 2022, Koskela, 2018, Koskela and Berenguer, 2019, Freund, 2021], and *(ii)* what sweepstakes reproduction may mean for the ecology and evolution of natural populations [Árnason, 2004, Hedrick, 2005, Hedgecock and Pudovkin, 2011, Waples, 2016, Eldon and Stephan, 2023, Árnason and Halldórsdóttir, 2015, Árnason et al., 2023, Eldon et al., 2016, Eldon, 2020, Vendrami et al., 2021].

Our results are relevant for inferring sweepstakes reproduction using data sampled from diploid populations. A diploid population is one where each individual carries a pair of chromosomes. For our purposes the population can be seen as consisting of pairs of (homologous) chromosomes. We will consider gene genealogies of a single contiguous non-recombining segment of a chromosome in a diploid panmictic population. The population evolves without selfing and according to sweepstakes reproduction. The evolution over a single generation is seen as occurring in two stages. In the first stage the current individuals (randomly paired) produce potential offspring according to a given law. In the second stage a given number of the potential offspring (conditional on there being enough of them) is then sampled uniformly and without replacement to survive to maturity and replace the current individuals; if the potential offspring are too few (fewer than the population size) the population remains unchanged over the generation.

A coalescent is a Markov process describing the random ancestral relations of sampled gene copies in a large population [Kingman, 1982c,a,b, Möhle and Sagitov, 2001, Pitman, 1999, Sagitov, 1999, Donnelly and Kurtz, 1999, Schweinsberg, 2000]. Multiple-merger coalescents describe gene genealogies in (arbitrarily large) populations evolving according to sweepstakes [Schweinsberg, 2003, Sagitov, 1999, Eldon and Wakeley, 2006, Sargsyan and Wakeley, 2008, Möhle and Sagitov, 2001, Huillet and Möhle, 2011, Der et al., 2012, Der and Plotkin, 2014, Gnedin et al., 2014, Matuszewski et al., 2018, Freund, 2020, Eldon, 2026]. A simultaneous multiple-merger coalescent (Xi-, Ξ-coalescent) is a coalescent where ancestral lineages may merge in two or more groups simultaneously [Schweinsberg, 2000, Möhle and Sagitov, 2001]. Diploidy intuitively induces simultaneous multiple mergers in gene genealogies from a diploid population evolving according to sweep-stakes [Möhle and Sagitov, 2003, Birkner et al., 2013a, 2018]. One should distinguish between haploidy and diploidy when it comes to comparing multiple-merger coalescents to data; Ξ-coalescents can predict patterns of genetic variation different from the ones admitting only asynchronous mergers [Birkner et al., 2013b, Blath et al., 2016, Spence et al., 2016, Eldon, 2026]. The coalescents derived here can be seen as mixtures of the Kingman coalescent [Kingman, 1982c,a] and specific families of Ξ-coalescents. Such mixtures also arise, for example, from models of strong positive selection [Durrett and Schweinsberg, 2005, Proposition 3.2].

Incorporating diploidy in studies of gene genealogies involves tracing ancestral lineages through homologous chromosomes of diploid individuals. Thus, a pair of ancestral lineages can be found in the same diploid individual without the lineages having merged. Viewed on the timescale applied when deriving a limit, such states occur over infinitesimally short periods of time, but prevent convergence in the *J*_1_-Skorokhod topology [Skorokhod, 1956]; one nevertheless has convergence in finite-dimensional distributions [Birkner et al., 2018]. A topology, which can be seen as an extension of the Skorokhod topology, has been proposed for convergence of Markov chains with such states [Landim, 2015]. However, we will follow Birkner et al. [2018] in proving convergence in the space of càdlàg paths for a process where the instantaneous states (occurring over an infinitesimal length of time in the limit of an arbitrarily large population) are simply ignored.

The layout of the paper: § 2 contains a brief background to coalescent processes, and to models of sweepstakes reproduction. In § 3 we state our main results in Theorems A, B, and C; § 4 contains numerical examples comparing functionals of gene genealogies and coalescents; § 6 holds proofs of Theorems A and B. Appendix A considers an example of a specific Ξ-coalescent; Appendices B–E contain brief descriptions of the simulation algorithms.

## 2 Background

For ease of reference we state notation used throughout.

### Definition 2.1

(Standard notation). *For any i* ∈ { 0, 1, 2,.. . }*write* ℕ_*i*_≡ { *i, i* + 1,.. . }, ℕ ≡ ℕ_1_. *Let N* ∈ ℕ *be fixed with* 2*N being the total number of diploid individuals (when the population size is constant)*.

*Asymptotic relations are seen to hold as the population becomes arbitrarily large, unless otherwise noted. ]*

*We let C, c, c*^′^,*K denote positive constants*.

*Write* [*n*] := {1, 2,…, *n*} *for any n* ∈ ℕ.

*Let ℰ*_*n*_ *denote the set of partitions on* [*n*] *for any n* ∈ ℕ *(ε*_1_ = {{{1}}}, *ε*_2_ = {{{1}, {2}}, {[2]}}, *etc.) We let* (*x*)_*m*_ *denote the falling factorial; for any real x, and any m* ∈ ℕ_0_,

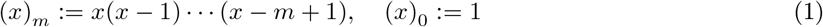

*For positive sequences* (*x*_*n*_)_*n*∈ℕ_ *and* (*y*_*n*_)_*n*∈ℕ_ *with* (*y*_*n*_) *bounded away from zero we will write*

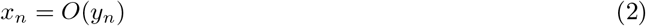

*if* lim sup_*n*→∞_ *x*_*n*_*/y*_*n*_ < ∞, *and*

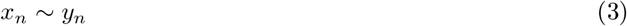

*if* lim_*n*→∞_ *x*_*n*_*/y*_*n*_ = 1. *We will also write*

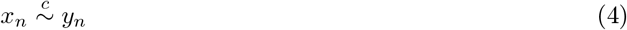

*if* lim_*n*→∞_ *x*_*n*_*/y*_*n*_ = *c for some (unspecified) constant c* > 0; (4) *is of course equivalent to x*_*n*_ *∼ cy*_*n*_. *Given x*_*n*_ *and y*_*n*_ *we will use* (4) *when we want to emphasize the conditions under which* (4) *holds*.

*Define*

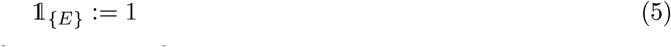

*when a given condition/event E holds, and take* 1_{*E*}_ = 0 *otherwise*.

*Take* |*A*| *to be the number of elements in a given finite set A*

*The abbreviation i.i.d. will stand for independent and identically distributed (random variables)*.

The introduction of the (annealed) coalescent [Hudson, 1983, Tajima, 1983, Kingman, 2000] marks a milestone in mathematical and empirical population genetics, as it is a rigorous probabilistic description of the random ancestral relations of sampled gene copies (chromosomes) [Kingman, 1982c,a,b, 1978]. A coalescent {*ξ*} ≡ {*ξ*(*t*); *t* ≥ 0} is a Markov chain (a Markov process with a countable state space and evolving in continuous time) taking values in the partitions of ℕ, such that the restriction {*ξ*^*n*^} ≡ {*ξ*^*n*^(*t*); *t* ≥ 0} to the partitions of [*n*] for a fixed *n* ∈ ℕ_2_ is still Markov and takes values in E_*n*_, the set of partitions of [*n*]. For a partition *ξ*^*n*^ ∈ E_*n*_ write 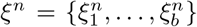, where 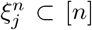, *b* = |*ξ*^*n*^| is the number of blocks in *ξ*^*n*^, 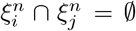 for *i* ≠ *j*, and 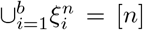. The only transitions are the merging of blocks of the current partition each time (we exclude elements such as recombination, population structure, or natural selection). Each block in a partition represents an ancestor to the elements (the leaves) in each of the blocks, in the sense that distinct leaves (corresponding to sampled gene copies) *i* and *j* are in the same block at time *t* 0 if and only if they share a common ancestor at time *t* in the past. At time zero, *ξ*^*n*^(0) = {{1},…, {*n*}}, and the time inf {*t* ≥ 0 : *ξ*^*n*^(*t*) = { [*n*] }}, where the partition { [*n*] } contains only the block [*n*], is when all the *n* leaves have found a common ancestor.

Here, we focus on diploid populations. Our aim is to describe the random ancestral relations of 2*n* gene copies from *n* randomly sampled diploid individuals. Following Birkner et al. [2018] and Möhle and Sagitov [2003] we define the state space

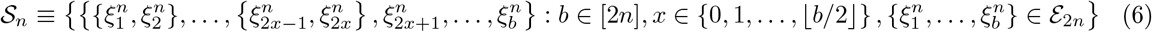

The elements of *S*_*n*_ when *x* = 0 (*x* is the number of diploid individuals carrying two ancestral blocks) are precisely the elements of ℰ_n_, so that ℰ_*n*_ ⊂ *S*_*n*_. The elements of _*n*_ corresponding to *x* > 0 have 2*x* blocks paired together in diploid individuals. A block in a partition *ξ*^*n*^ ∈ *S*_*n*_ is ‘ancestral’ in the sense that it is an ancestor of the sampled gene copies (arbitrarily labelled) whose labels are contained in the block. Let {*ξ*^*n,N*^ } ≡ {*ξ*^*n,N*^ (*τ* ); *τ* ∈ ℕ_0_}be a Markov sequence (a Markov process with a countable state space and evolving in discrete time) with values in _*n*_. We will refer to {*ξ*^*n,N*^} as the *annealed ancestral process*; leaves (sampled gene copies) labelled *i* and *j* are in the same block at time *τ* if and only if they share a common ancestor at time *τ* in the past. We measure time going from the present into the past, we take *ξ*^*n,N*^ (0) = {{1, 2},…, {2*n* 1, 2*n* }}, and the only transitions are the merging of blocks of the current partition (we exclude further elements such as recombination or population structure). We will also refer to the process keeping track of the block sizes of a given ancestral process resp. coalescent as an ancestral process resp. coalescent.

A central quantity in deriving limits of {*ξ*^*n,N*^ }is the coalescence probability [Sagitov, 1999].

### Definition 2.2

(The coalescence probability). *Define c*_*N*_ *as the probability that two given gene copies in separate diploid individuals from a given generation derive from the same parent gene copy*

We will show that *c*_*N*_ → 0, and that 1*/c*_*N*_ is proportional to (at least) *N/* log *N* for the models we will consider. It holds that 1*/c*_*N*_ is the correct scaling of time of the ancestral process for proving convergence [Sagitov, 2003, Equation 1.4]. If *c*_*N*_ → 0, the limiting coalescent evolves in continuous time, with one unit of time corresponding to 1*/c*_*N*_ generations [Schweinsberg, 2003, Möhle and Sagitov, 2001].

For each *N* ∈ ℕ let *v*_1_,…, *v*_*N*_ denote the random number of surviving offspring from the *N* current parent pairs at some arbitrary time. We will assume that (*v*_1_,…, *v*_*N*_ ) are independent across generations (and identically distributed when the population size is constant). Definition 2.2 then gives, with 2*N* being the population size (recall the notation from Definition 2.1),

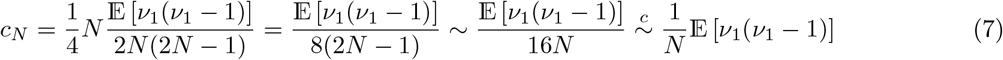

### 2.1 The population model

In this section we define more precisely the evolution of a diploid population evolving without selfing.

#### Definition 2.3

(Evolution of a diploid population). *Consider a diploid panmictic population. In each generation the current individuals arbitrarily form parent pairs, each pair is made up of distinct individuals, each individual shows up in at most 1 pair (individuals are monogamous), and selfing is prohibited. The parent pairs thus formed independently produce random numbers of diploid potential offspring according to some given law. Each offspring receives two chromosomes (gene copy), one chromosome from each of its two parents, with each inherited chromosome sampled independently and uniformly at random from among the two parent chromosomes. If the total number of potential offspring is at least some given number (*2*N when the population size is constant), we sample the given number of the potential offspring uniformly at random without replacement to survive to maturity and replace the current individuals; otherwise we assume the population is unchanged over the generation (all the potential offspring perish before reaching maturity)*.

#### Remark 2.4

(Illustrating Definition 2.3). *The mechanism described in Definition 2.3 is illustrated below, where* {*a, b*} *denotes a diploid individual carrying gene copies a, b, and* {{*a, b*}, {*c, d*}} *denotes a pair of diploid individuals (a parent pair). Here we have arbitrarily labelled the gene copies in order to illustrate the evolution over one generation. In this example the population size stays constant at* 2*N diploid individuals*.

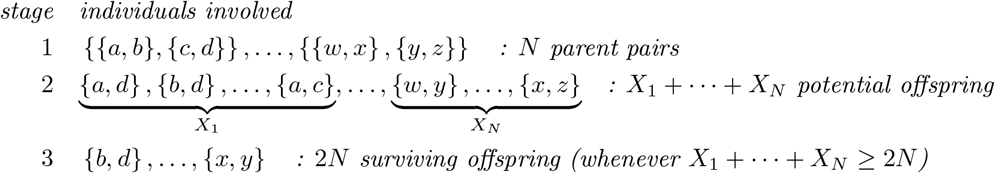

*In stage 1 above, the current* 2*N diploid individuals randomly form N pairs; in stage 2 the N pairs formed in stage 1 independently produce random numbers X*_1_,…, *X*_*N*_ *of potential offspring, where each diploid offspring receives one gene copy (sampled uniformly at random) from each of its two parents; in the third stage* 2*N of the X*_1_ +…+*X*_*N*_ *potential offspring (conditional on there being at least* 2*N of them) are sampled uniformly and without replacement to survive to maturity and replace the parents*.

#### Remark 2.5

(Relation to Birkner et al. [2018]). *The reproduction mechanism described in Definition 2.3 is a special case of the one studied by Birkner et al. [2018]. They consider an array* 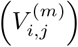 *of exchangeable offspring numbers where* 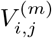 *is the random number of* surviving *offspring produced in generation m by individuals i and j (arbitrarily labelled, and with* 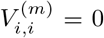). *The idea is that any individual may produce offspring with more than one individual in the same generation (promiscuous mating). In principle it should be possible to use the framework of Birkner et al. [2018]. However, as we are interested in comparing a given ancestral process to the limiting coalescent, we follow a simpler framework where in any given generation each diploid individual produces offspring in collaboration with exactly one other individual (monogamous mating). We elect to leave an extension of [Birkner et al*., *2018] to random environments to future work. A* Ξ*-coalescent with an atom at zero is briefly discussed in [Birkner et al*., *2018, Section 1.1]. Here, we specifically consider scenarios where the limiting* Ξ*-coalescent does have an atom at zero. In our case the time-scaling used for viewing gene genealogies in a large population is affected. The concept of time-scaling is relevant for explaining genetic variation (see § 5)*.

Due to diploidy (see Definition 2.3 and Remark 2.4) the ancestral process may enter states where two gene copies (ancestral to the sampled ones) reside in the same diploid individual. This is why in Definition 2.2 we require the two gene copies to be in distinct diploid individuals. It should also be clear from the illustration in Remark 2.4 why one could expect to see simultaneous mergers in the genealogy of a sample from a diploid population where large families regularly occur. For family *i* we sample 2*X*_*i*_ times (with replacement) from the 4 parental chromosomes involved. Then, when a parent pair produces of order *O*(*N* ) number of offspring, we can have up to 4 groups of offspring chromosomes of order *O*(*N* ) belonging to the same tetrad (quadruplet) of parent chromosomes, which translates to simultaneous mergers of ancestral lineages.

The following population model was initially introduced for a haploid population. Suppose *α, C* > 0 are fixed and *X* is the random number of potential offspring produced by an arbitrary individual (gene copy), and that [Schweinsberg, 2003, Equation 11]

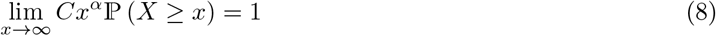

From the pool of potential offspring produced at the same time *N* of them (conditional on there being at least *N* potential offspring) are sampled uniformly at random and without replacement to survive to maturity and replace the parents (recall Definition 2.3). When 1 ≤ *α* < 2 (8) gives rise to the (asynchronous) Beta(2 −*α, α*)-coalescent [Schweinsberg, 2003, Theorem 4c], a family of multiple-merger coalescents studied to some extent [Birkner et al., 2005, Dahmer et al., 2014, Berestycki et al., 2007, 2008, Kersting et al., 2014, Birkner et al., 2024]. We will consider extensions of (8) applied to diploid populations evolving as described in Definition 2.3.

### 2.2 Simultaneous multiple-merger coalescents

We will be concerned with simultaneous multiple-merger coalescents (Ξ-coalescents,Xi-coalescents) where ancestral lineages may merge in several groups simultaneously [Schweinsberg, 2000, Möhle and Sagitov, 2001, Sagitov, 2003, Möhle and Sagitov, 2003]. Sweepstakes reproduction, recurrent strong bottlenecks, or strong positive selection, are examples of mechanisms generating Ξ-coalescents [Sargsyan and Wakeley, 2008, Eldon, 2026, Birkner et al., 2018, 2013a, Möhle and Sagitov, 2003, Birkner et al., 2009, Durrett and Schweinsberg, 2005, Schweinsberg and Durrett, 2005, Taylor and Véber, 2009].

Define

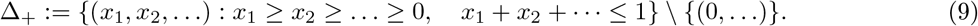

Suppose the merging measure Ξ of a Ξ-coalescent on Δ_+_ ∪ {(0,.. .)} is of the form (*c, c*^′^ ≥ 0 fixed)

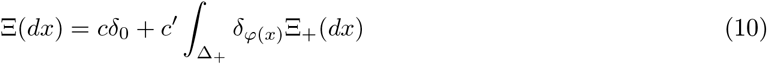

with Ξ_+_ a finite measure on Δ_+_, and *φ* : Δ_+_ → Δ_+_ a given map. Then, with *n* ≥ 2 active blocks in the current partition, *k*_1_,…, *k*_*r*_ ≥ 2, *r* ∈ ℕ, and 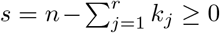, the rate at which *k*_1_ +… + *k*_*r*_ ∈ {2,…, *n*} blocks merge in *r* groups with group *j* of size *k*_*j*_ is given by (*c, c*^′^ ≥ 0 fixed, and *ϕ* the identity map)

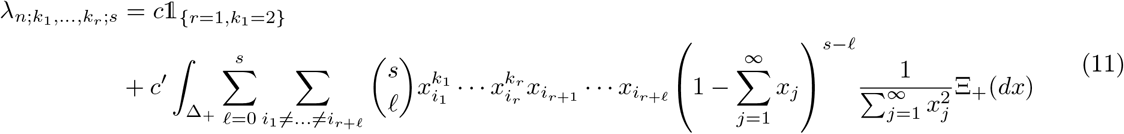

[Schweinsberg, 2000, Möhle and Sagitov, 2001]. The constant *c* in (11) is the mass the merging measure Ξ assigns to {(0,.. .)}, and *c*^′^ the mass assigned to Δ_+_. The application of (8) to a diploid panmictic population of constant size evolving according to Definition 2.3 leads to a Ξ-coalescent without an atom at zero (corresponding to *c* = {Ξ (0,.. .)} = 0 in (11)) and driving measure Ξ_+_ of the form

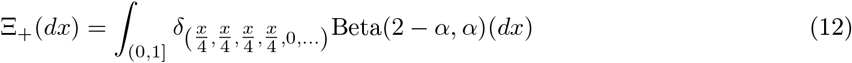

where Beta(2 − *α, α*) stands for the beta density with parameters 2 − *α* and *α* for 1 < *α* < 2 [Birkner et al., 2018, Proposition 2.5(2); Equation 29]. The form of Ξ_+_ in (12) says that the measure Ξ_+_ is of the form Ξ_+_ = Beta(2 -*α, α*) ◯ φ^−1^, where φ : (0, 1] → Δ_+_ takes values φ(*x*) ≡ (*x/*4, *x/*4, *x/*4, *x/*4, 0,…).

The *x* in (12) can be seen as the fraction of surviving offspring produced by an arbitrary parent pair (a pair of pairs of gene copies). The ancestral lines belonging to the same family are then split (uniformly at random and with replacement) among the four parental chromosomes, and the ones assigned to the same chromosome are merged. We will consider extensions of (8) that, when combining the results, allow us to take 0 < *α* < 2. Moreover, we will consider a truncated (incomplete) version of the Beta(2 − *α, α*)-coalescent.

When 0 < *α* < 1 one obtains, using (8) for a haploid panmictic population of constant size, a discrete-time Ξ-coalescent associated with the Poisson-Dirichlet distribution with parameter (*α*, 0) [Schweinsberg, 2003, Theorem 4(d)]. The Poisson-Dirichlet distribution [Kingman, 1975] has found wide applicability, including in population genetics [Feng, 2010, Bertoin, 2006, Sagitov, 2003, Eldon, 2026]. We will be concerned with the two-parameter Poisson-Dirichlet(*α, θ*) distribution for 0 < *α* < 1 and *θ* > −*α* restricted to *θ* = 0.

#### Definition 2.6

(A Poisson-Dirichlet(*α*, 0)-coalescent; [Schweinsberg, 2003]). *Let* 0 < *α* < 1 *be fixed, recall the simplex* Δ_+_ *in* (9), *write x* = (*x*_1_, *x*_2_,.. .) *for x* ∈ Δ_+_, *and* 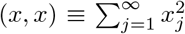. *Let F*_*α*_ *be a probability measure on* Δ_+_ *associated with the Poisson-Dirichlet*(*α*, 0)*-distribution, and* Ξ_*α*_ *a measure on* Δ_+_ *given by*

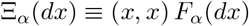

*A Poisson-Dirichlet*(*α*, 0)*-coalescent is a discrete-time* Ξ*-coalescent with* Ξ*-measure* Ξ_*α*_ *and no atom at zero. The transition probability of merging blocks in r groups of size k*_1_,…, *k*_*r*_ ≥ 2 *with current number of blocks b* ≥ *k*_1_ + … + *k*_*r*_, *and s* ≡ *b* − *k*_1_ −· · · − *k*_*r*_, *is given by (recall* (1)),

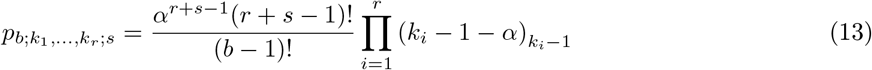

(cf. [Schweinsberg, 2003, Equation 13], [Eldon, 2026, Equation 12]). The Ξ_+_-measure (recall (11)) of the Poisson-Dirichlet(*α*, 0)-coalescent is of the form

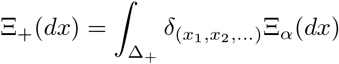

### 2.3 Two Ω-Ξ -coalescents

We define two Ξ-coalescents, the Ω-*δ*_0_-Beta(*γ*, 2 −*α, α*)-coalescent, and the Ω-*δ*_0_-Poisson-Dirichlet(*α*, 0)-coalescent (see Appendix A for the Ω-Beta(2 − *β, β*)-Poisson-Dirichlet(*α*, 0)-coalescent) that we will need for stating our results. We use the prefix Ω to denote a Ξ-coalescent where the ancestral lines involved in each group of mergers are assigned (uniformly at random and with replacement) into four subgroups, and the lines assigned to the same subgroup are merged; an Ω-Ξ-coalescent is a Ξ-coalescents with the merging measure Ξ as given in (10), with

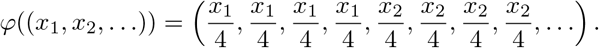

#### Definition 2.7

(The Ω-*δ*_0_-Beta(*γ*, 2 −*α, α*)-coalescent). *Let* **k** ≡ (*k*_1_,…, *k*_*r*_) *for r* ∈ [4], *where* 2 ≤ *k*_*i*_ ≤ *n, k* ≡ *k*_1_ +…+ *k*_*r*_ ≤ *n, and s* ≡ *n* −*k for n* ≥ 2. *The* Ω*-δ*_0_*-Beta*(*γ*, 2 −*α, α*)*-coalescent, for* 0 < *γ* ≤1 *and* 0 < *α* < 2, *is a continuous-time* Ξ*-coalescent (recall* (11)*) on the partitions of* [*n*], *where the only transitions are the merging of blocks of the current partition. The transition rate, at which r* ∈ [4] *(simultaneous when r* ≥ 2*) mergers occur with the merger sizes given by k*_1_,…, *k*_*r*_ *and* 2 ≤ _*i*_ *k*_*i*_ ≤ *n, is (c, c*^′^ > 0 *fixed)*

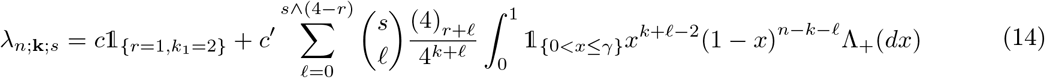

*The finite measure* Λ_+_ (14) *on* (0, 1] *is given by, for* 0 < *γ* ≤ 1 *and* 0 < *α* < 2 *both fixed*,

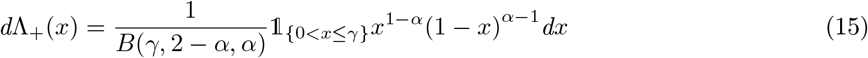

*In* (15) *B* 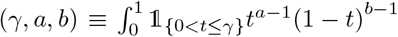 *dt for a, b* > 0, *and* 0 < *γ* ≤ 1, *is the beta function. Then, λ*_*n*;**k**;*s*_ *in* (14) *takes the form (x ∧ y is the smaller of x and y)*

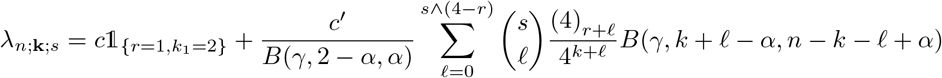

To see that the sum in (14) follows from (11) when Ξ_+_ is as given, recall that {*ξ*^*n*^} is restricted to ℰ_*n*_ (recall Definition 2.1). It then suffices to see that given *k* ancestral lines merging in *r* groups, we can have up to *s ∧* (4 − *r*) additional lines each assigned to one ‘free’ parental chromosome (to which none of the *k* lines are assigned) and there are (4)_*r*+ *ℓ*_ equivalent ways of ordering the *r* +*l* parent chromosomes receiving an ancestral line [Birkner et al., 2013a, Equation 27]. The merging measure of the Ω-*δ*_0_-Beta(*γ*, 2 − *α, α*)-coalescent defined in Definition 2.7 is of the form (recall (12))

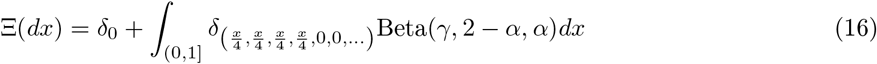

where Beta(*γ*, 2 −*α, α*) is the law on (0, 1] with density (15).

The *δ*_0_-Poisson-Dirichlet(*α*, 0)-coalescent is a continuous-time Poisson-Dirichlet(*α*, 0)-coalescent [Schweinsberg, 2003] with an atom at zero; it describes gene genealogies in haploid populations evolving in a random environment similar to the ones we will consider [Eldon, 2026].

#### Definition 2.8

(The *δ*_0_-Poisson-Dirichlet(*α*, 0)-coalescent). *The δ*_0_*-Poisson-Dirichlet*(*α*, 0)*-coalescent is a continuous-time* Ξ *coalescent (recall* (11)*) taking values in ℰ*_*n*_ *with* Ξ*-measure* Ξ = *δ*_0_ + Ξ_+_, *where* Ξ_+_ = Ξ_*α*_ *and* Ξ_*α*_ *is as in Definition 2.6. The only transitions are the merging of blocks of the current partition. Given ℰn blocks in a partition, k*_1_,…, *k*_*r*_ ≥ 2 *with* 2 ≤ *k*_1_ +… + *k*_*r*_ ≤ *n denoting the merger sizes of r (simultaneous when r* ≥ 2*) mergers, s* = *n* − *k*_1_ − … − *k*_*r*_, *a* (*k*_1_,…, *k*_*r*_)*-merger occurs at rate (where c, c*^′^ > 0 *fixed)*

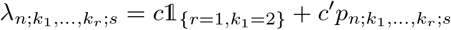

*where* 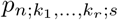 *is as in* (13).

#### Definition 2.9

(A Ω-*δ*_0_-Poisson-Dirichlet(*α*,0)-coalescent). *A* Ω*-δ*_0_*-Poisson-Dirichlet*(*α*, 0)*-coalescent is a δ*_0_*-Poisson-Dirichlet*(*α*, 0)*-coalescent (recall Definition 2.8) where the blocks in each group of merging blocks are split among four subgroups independently and uniformly at random and with replacement, and the blocks assigned to the same subgroup are merged*.

The merging measure of the Ω-*δ*_0_-Poisson-Dirichlet(*α*, 0)-coalescent in Definition 2.9 is given by

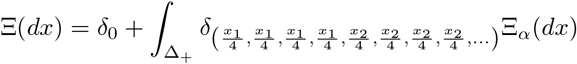

where Ξ_*α*_ is as in Definition 2.6 and Δ_+_ as in (9).

#### Remark 2.10

(Extending [Koskela and Berenguer, 2019]). *The* Ω*-δ*_0_*-Poisson-Dirichlet*(*α*, 0)*-coalescent extends the coalescents considered by Koskela and Berenguer [2019], who consider coalescents based on population models where at most one large family (with an offsping number proportional to the population size) occurs with non-negligible probability at any given time in a large population [Koskela and Berenguer, 2019, Equations 9 and 10], thus leading to* Ξ*-coalescents with measure of the form as in* (12) *or* (16). *In our model behind the* Ω*-δ*_0_*-Poisson-Dirichlet*(*α*, 0)*-coalescent, when using* (25) *with a* = *α* ∈ (0, 1), *we expect to see of order* O(*N* ^1−*α*^) *families with at least N potential offspring. We expect, during the history of a sample, to use α on the order of* O(1) *number of times for sampling numbers of offspring*.

### 2.4 Equivalent convergence conditions

Equivalent conditions for convergence of { ξ^*n,N*^} to Ξ -coalescents are summarised by Birkner et al. [2018]. Convergence, in the sense of convergence of finite-dimensional distributions of {*ξ*^*n,N*^} to a Ξ-coalescent, depend on the existence of the limits (recall *c*_*N*_ from Definition 2.2 and (7))

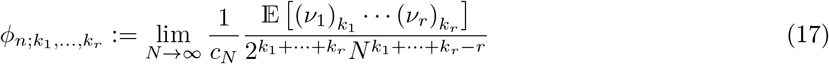

for all *r* ∈ ℕ, *k*_1_,…, *k*_*r*_ ≥2 are the merger sizes, 2 ≤*k*_1_ + …+ *k*_*r*_ ≤*n*, and *n* is the current number of blocks [Möhle and Sagitov, 2001, Birkner et al., 2018]. See [Birkner et al., 2018, Theorem A.5] for a summary of equivalent conditions for convergence to a Ξ-coalescent. Existence of the limits in (17) is equivalent to

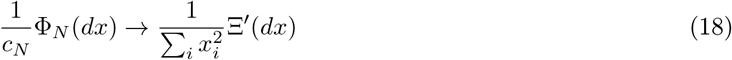

vaguely on Δ_+_ as ℕ → ∞; Ξ^′^ is a probability measure on Δ_+_, and

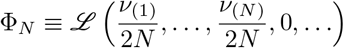

is the law of the ranked individual-wise offspring frequencies *ν*_(1)_*/*(2*N* ) ≥ *ν*_(2)_*/*(2*N* ) ≥… ≥*ν*_(*N*)_*/*(2*N* ). Write

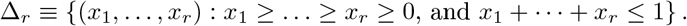

Let *F*_*r*_ be a symmetric measure on Δ_*r*_ for *r* ∈ ℕ. Equivalent to equivalent conditions (17) and (18) are the two conditions [Birkner et al., 2018, Condition II in Appendix A] (see also [Möhle and Sagitov, 2001, Equations 21 and 22])

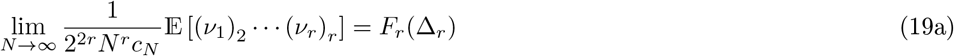

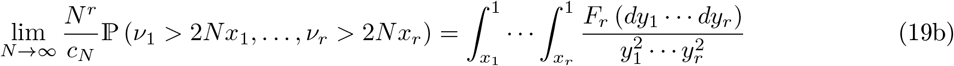

with the limits in (19a) holding for all *r* ∈ ℕ, and (19b) holding for points of continuity for *F*_*r*_.

## 3 Mathematical results

In § 3.1 we present the model behind the main results. Theorems A and B record the results for constant population size (§ 3.2), and Theorem C for (deterministically) varying population size (§ 3.3).

### 3.1 The law for the number of potential offspring

In this section we present the law for the random number of potential offspring underlying the main results. For ease of reference we first state key notation.

#### Definition 3.1.

*(Notation) For each N* ∈ ℕ, *we let X*_1_,…, *X*_*N*_, *resp. ν*_1_,…, *ν*_*N*_, *denote the random number of potential resp. surviving offspring produced in an arbitrary generation by the current N parent pairs (arbitrarily formed; recall Definition 2.3). Recall that* 2*N is the population size, the number of diploid individuals (pairs of gene copies) at any time when the population size is constant, and then ν*_1_+·· ·+*ν*_*N*_ = 2*N for each N* ∈ ℕ. *The X*_1_,…, *X*_*N*_ *are independent. Write, for each N* ≥ 2,

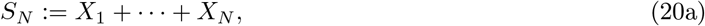

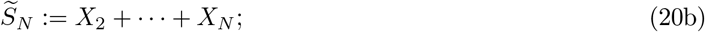

*S*_*N*_ *is the total number of diploid potential offspring produced in a given generation. Write*

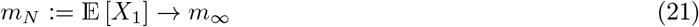

*for the expected value of X*_1_, *with m*_∞_ *denoting the (assumed fixed) mean as N* → ∞ *( we will identify conditions for m*_∞_ *to be finite; see Lemmas 6.7 and 6.14)*.

*Write, with ζ*(*N* ) *a positive function of N (see* (25)),

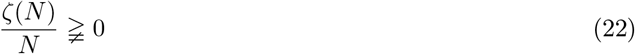

*if either ζ*(*N* )*/N converges to a strictly positive constant, or ζ*(*N* )*/N diverges, as N* → ∞.

*We write, with κ* ≥ 2 *a given positive constant (recalling* 1_{·}_ *from* (5)),

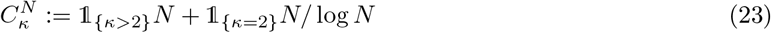

Now we state the distribution for the number of potential offspring underlying our results. Suppose *X* represents the random number of potential offspring produced by an arbitrary pair of diploid individuals (parent pair). We write, with *a* > 0 fixed, and *ζ*(*N* ) a positive deterministic function of *N*, with *a* and *ζ*(*N* ) as given in each case,

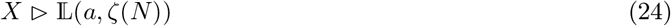

when the probability mass function of the law of *X* is bounded by, for *k* ∈ {2, 3,…,*ζ*(*N* )} and each *N* ∈ ℕ,

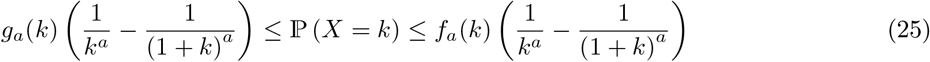

We assign the remaining mass (outside {2, 3,…,*ζ*(*N* ) }) to {*X* ∈ {0, 1}} ; how the mass is distributed between 0 and 1 will become irrelevant in the limit (see Lemma 6.6). We take {*f*_*a*_}, {*g*_*a*_} to be families of bounded positive functions on ℕ such that p (*X* ≤ *ζ*(*N* )) = 1 for any *a* > 0 and *N* ∈ ℕ. We will identify conditions on *g*_*a*_ and *f*_*a*_ such that the ancestral process converges to a non-trivial limit. The law (25) is an extension of the one in (8); with *X* distributed as in (8) we have 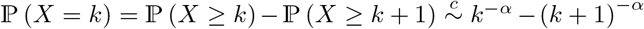 as *k* → ∞. The bounds on the mass function in (25) are the same for each *N* ∈ ℕ. We can equivalently define *X*^(*N*)^ ≡ *X*1_{*X*∈[*ζ*(*N*)]}_ + 1_{*X*>*ζ*(*N*)}_, and take *X*_1_,…, *X*_*N*_ to be independent copies of *X*^(*N*)^ (see Remark 3.2) for each *N* ≥ 2. See [Chetwynd-Diggle and Eldon, 2026, Eldon, 2026] for (25) in the context of haploid populations.

#### Remark 3.2

(A sequence of independent random variables). *The law* (25) *is seen to hold for the random numbers X*_1_,…, *X*_*N*_ *of potential offspring of the current parent pairs (N pairs when the population size is* 2*N) for each N* ∈ ℕ. *Equivalently, one may consider a sequence* 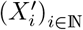 *of independent random variables, where* 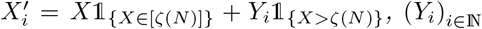 *is a sequence of independent random variables suitably defined, and X* ⊳ L (*a, ζ*(*N* )) *(recall* (24)*). We will work with such sequences when we prove almost sure convergence of S*_*N*_ */*(*Nm*_*N*_ ) *(recall* (20a) *and* (21) *in § 3.1)*.

We assume *g*_a_ (*k*) ≤ *f*_a_ (*k*) for all *k*. We define, for any 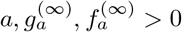 fixed,

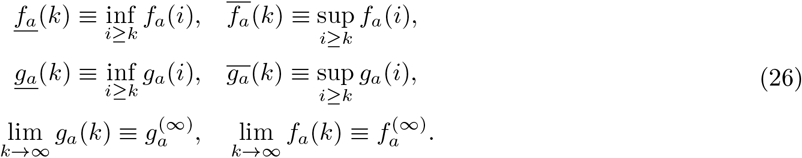

We then have, for *k* ∈ {2, 3,…,*ζ*(*N* )}, with the law of *X* as in (25) for each *N* ≥ 2,

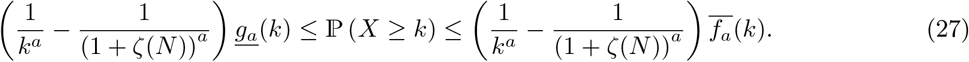

The functions *f*_*a*_ and *g*_*a*_ in (25) can be chosen such that ℙ (*X* = *k* − 1) ≥ ℙ (*X* = *k*) for *k* ∈ { 3,…,*ζ*(*N* ) }, and that E [*X*] > 2 for all *N* large enough. The monotonicity requirement is reasonable (but not necessary). The requirement that E [*X*_*i*_] > 2 for all *i* ∈ [*N* ] and *N* ∈ ℕ results in ℙ (*X*_1_ +…+ *X*_*N*_ < 2*N* ) decreasing exponentially in *N* (Lemma 6.6), where *X*_1_,…, *X*_*N*_ are as in Definition 3.1 (recall Remark 3.2).

#### Remark 3.3

(Assumption on *g*_*a*_ and *f*_*a*_ in (25)). *With X* l⊳ L (*a, ζ*(*N* )) *(recall* (24)*) we will assume that the functions f*_*a*_ *and g*_*a*_ *in* (25) *are such that* E [*X*] > 2 *for all N* ∈ ℕ *(recall that parent pairs produce offspring), and that the constant m*_∞_ > 2 *(recall* (21)).

### 3.2 Constant population size

In this section we define the random environments, and state the convergence results for constant population size. Results for when the population size varies in time are stated in § 3.3.

The coalescent in Definitions 2.7 and 2.9 arises in specific cases when the population evolves as follows.

#### Definition 3.4

(A random environment). *Suppose a diploid panmictic population evolves as in Definition 2.3. Fix* 0 < *α* < 2, *and* 2 ≤ *κ. Recall X*_1_,…, *X*_*N*_ *from Definition 3.1. Write E for the event when X*_*i*_ ⊳ L(*α, ζ*(*N* )) *for all i* ∈ [*N* ] *(recall* (24)*), and E*^c^ *for the event when κ replaces α in E (X*_*i*_ ⊳ L(*κ, ζ*(*N* )) *for all i* ∈ [*N* ]*). Let* 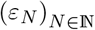 *be a positive sequence with* 0 < *ε*_*N*_ < 1 *for all N. It may hold that ε*_*N*_ → 0 *as N* → ∞. *Take, for each N* ∈ ℕ,

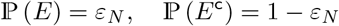

Definition 3.4 says that the *X*_1_,…, *X*_*N*_ are i.i.d. copies of *X* where 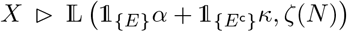 . The idea represented by Definition 3.4 is that most of the time (with probability 1 − *ε*_*N*_ ) small (relative to the population size) numbers of offspring per family are produced with high probability through *κ* (event *E*^c^); occasionally, however, (with probability *ε*_*N*_ ) environmental conditions are favourable for producing an increased number of offspring through *α* (event *E*), and then the current parent pairs will all (independently) produce a random number of potential offspring using (25) with *a* = *α*.

The coalescents resulting from Definition 3.4 are described in Theorem A; § 6.1 contains a proof of Theorem A.

#### Theorem A

(Coalescents under Definition 3.4). *Suppose a diploid population evolves according to Definitions 2.3 and 3.4, and that Assumption 3.3 holds. Then* {*ξ*^*n,N*^ ( *t/c*_*N*_ ); *t* ≥ 0} *converges, as N* → ∞, *in the sense of convergence of finite-dimensional distributions, to a conti nuous-time coalescent* { *ξ*^*n*^} ≡ {*ξ*^*n*^(*t*); *t* ≥ 0} *as specified in each case. When* 1 ≤ *α* < 2 *we take ε*_*N*_ = *cN*^*α*−2^ 1_{*κ*>2}_ + 1_{*κ*=2}_ log *N with c* > 0 *fixed*.

1. *The limiting coalescent* {*ξ*^*n*^} *is the Kingman coalescent when one of*
  a. 0 < *α* < 1, *ζ*(*N* )^3−*α*^*/N* ^2^ → 0 *as N* → ∞, *and* lim sup _*N* →∞_ *ε*_*N*_ */c*_*N*_ < ∞;
  b. 1 ≤ *α* < 2, *ε*_*N*_ *is as stated above, and ζ*(*N* )*/N* → 0 *as N* → ∞; *is in force*.
2. *Suppose* 1 ≤ *α* < 2 *and ζ*(*N* )*/N ≩* 0 *(recall* (22) *in Definition 3.1). Then* {*ξ*^*n*^} *is the* Ω*-δ*_0_*-Beta*(*γ*, 2 − *α, α*)*-coalescent as in Definition 2.7. Given n blocks the transition rate for a (simultaneous multiple) merger with group sizes k*_1_,…, *k*_*r*_ *for r* ∈ [4] *is (recall m*_∞_ *from* (21))

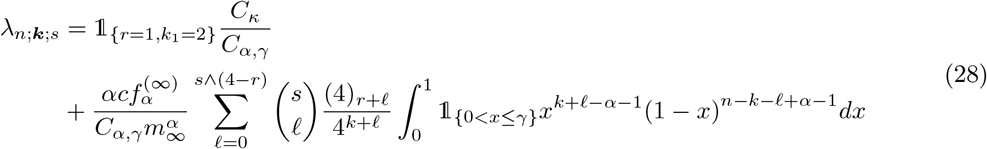

*where we have, recalling B*(*γ*, 2 − *α, α*) *from Definition 2.7, and* 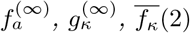 *from* (26)

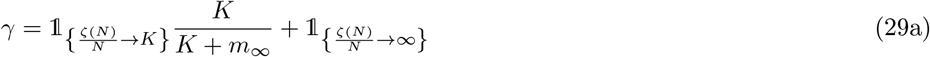

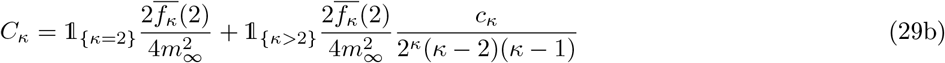

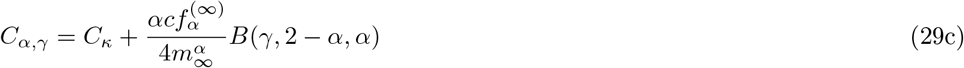

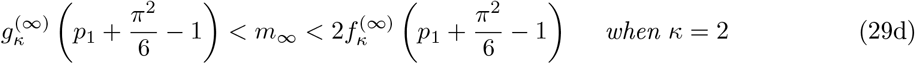

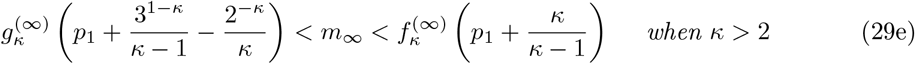

*where in* (29d) *and* (29e) *p*_1_ = ℙ (*X* = 1) *with X* ⊳ L(*κ, ζ*(*N* )) *(recall* (24)*). Moreover, it holds that κ* +2 < *c*_*κ*_ < *κ*^2^ (29b) *when κ* > 2, *and λ*_2;2;0_ = 1. *Recall that ζ*(*N* )*/N ≩* 0 *means (by assumption) that either ζ*(*N* )*/N converges to a strictly positive constant, or that ζ*(*N* )*/N diverges, as reflected in* (29a).
3. *Suppose* 1*/*2 < *α* < 1, *and ζ*(*N* )*/N* ^1*/α*^ → ∞ . *Then* {*ξ*^*n*^} *is the* Ω*-δ*_0_*-Poisson-Dirichlet*(*α*, 0)*-coalescent as in Definition 2.9. A transition proceeds by sampling group sizes using the δ*_0_*-Poisson-Dirichlet*(*α*, 0)*-coalescent, the blocks (ancestral lineages) in each group are split among 4 subgroups independently and uniformly at random, and the blocks assigned to the same subgroup are merged. When there are n blocks, the total rate of observing a (simultaneous multiple) merger, with group sizes k*_1_,…, *k*_*r*_ ≥ 2 *such that ∑*_*i*_ *k*_*i*_ ≤ *n, is*

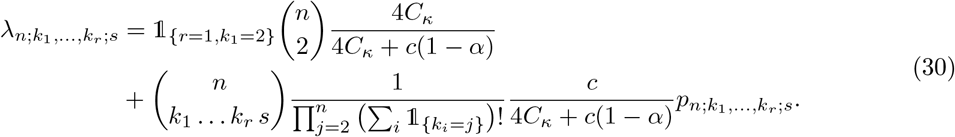

*In* (30), *C*_*κ*_ *is as in* (29b), *m*_∞_ *as in* (29d) *or* (29e), 2 ≤ *k*_1_,…, *k*_*r*_ ≤ *n, ∑*_*i*_ *k*_*i*_ ≤ *n, and s* = *n* − ∑_*i*_ *k*_*i*_. *The positive constant c in* (30) *is such that* 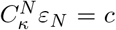 *for all N* ∈ ℕ. *In* (30) 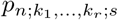*is as in* (13).

*For all the cases above we have, as N* → ∞ *(recall* (4)),

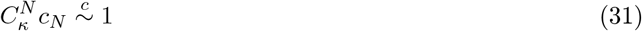

*with c*_*N*_ *as in Definition 2.2 (recall* (7)); 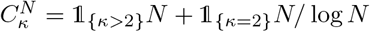 *as in* (23).

#### Remark 3.5

(The range of *α* in Case 3 of Theorem A). *The proof of Case 3 of Theorem A shows that we require ζ*(*N* )*/N* ^1*/α*^ → ∞. *Suppose ζ*(*N* ) = *N*^*δ*+1*/α*^ *for some fixed* 0 < *δ* ≪ 1. *Assuming* 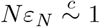 *forces us to take* 1*/*2 < *α* < 1 *to guarantee that m*_∞_ < ∞ *by Lemma 6.7. In Case 3 of Theorem A, the bound ζ*(*N* ) *is such that ζ*(*N* )*/N* ^1*/α*^ → ∞, *and seen to be at most of the form N*^*δ*+1*/α*^.

#### Remark 3.6.

*Taking ζ*(*N* ) = *N*^*x*^ *for some fixed x* > 0 *we see that the condition ζ*(*N* )^3−*α*^*/N* ^2^ → 0 *in Case 1 of Theorem A corresponds to x* < 2*/*(3 − *α*) < 1 *when* 0 < *α* < 1, *which is a stronger ask of ζ*(*N* ) *than the requirement ζ*(*N* )*/N* → 0 *when* 1 ≤ *α* < 2.

#### Remark 3.7

(The parameters of the Ω-*δ*_0_-Beta(*γ*, 2 − *α, α*)-coalescent). *The transition rates of the* Ω*-δ*_0_*-Beta*(*γ*, 2 − *α, α*)*-coalescent are functions of the parameters α, γ, c, and κ, and also of the mean m*_∞_.

#### Remark 3.8

(Poisson-Dirichlet-coalescents and stick-breaking constructions). *The Poisson-Dirichlet*(*α*, 0)*-coalescent can be sampled using a stick-breaking construction [Feng, 2010]. Similar constructions feature in* Ξ*-coalescents generated by recurrent selective sweeps [Durrett and Schweinsberg, 2005, Section 3]*.

As detailed in [Birkner et al., 2018, § 1], and formalised in [Birkner et al., 2018, Corollary 1.2], by tracking only completely dispersed states we have weak convergence on *D*([0, ∞), *ℰ*_2*n*_) (the set of ℰ_2*n*_ valued càdlàg paths with Skorokhod’s *J*_1_ topology [Ethier and Kurtz, 2005, Chapter 3.4]). The proof of Corollary 3.9 is identical to the one of [Birkner et al., 2018, Corollary 1.2]. For *ξ*^*n*^ ∈ *S*_*n*_ (recall (6)) define the map `cd` : *S*_*n*_ → ℰ_2*n*_ (with cd standing for ‘complete dispersal’) by

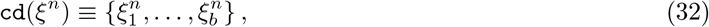

where *b* is the number of blocks in *ξ*^*n*^ (recall (6)). Then *ξ*^*n*^ = cd(*ξ*^*n*^) for all *ξ*^*n*^ ∈ ℰ_2*n*_.

#### Corollary 3.9

(Weak convergence of {*ξ* ^*n,N*^} on *D*([0, ∞), ℰ_2*n*_)). *Let* 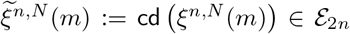 *for all m. Under the conditions of Theorem A* 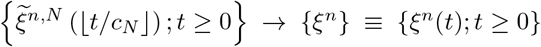 *weakly on D* ([0, ∞), ℰ_2*n*_) *as N* → ∞, *with* {*ξ*^*n*^} *as given in Theorem A*.

We will also consider an environment where the *X*_1_,…, *X*_*N*_ for all *N* stay independent but may not always be identically distributed.

#### Definition 3.10

(A random environment). *Suppose a diploid population evolves as in Definition 2.3. Fix* 0 < *α* < 2, *and κ* ≥ 2. *Recall X*_1_,…, *X*_*N*_ *from Definition 3.1. Write E*_1_ *for the event, when there exists exactly one i* ∈ [*N*] *where X*_*i*_ ⊳ L(*α, ζ*(*N* )), *and X*_*j*_ ⊳ L(*κ, ζ*(*N* )) *for all j* ∈ [*N* ] \ {*i*}, *and for each N* ∈ ℕ. *When E*_1_ *occurs the index i is picked uniformly at random. Write* 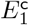 *when κ replaces α in E*_1_; *when* 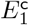 *occurs the X*_1_,…, *X*_*N*_ *are i.i.d. copies of X where X* ⊳ L(*κ, ζ*(*N* )), *for each N* ∈ ℕ. *Let* 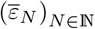 *be a positive sequence with* 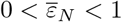 *for all N. It may hold that* 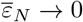 *as N* → ∞. *Suppose, for all N*,

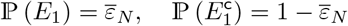

Definition 3.10 says, that in each generation there exists an *I* ∈ [*N* ] where 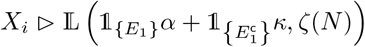, and *X*_*j*_ ⊳ L (*κ, ζ*(*N* )) for all *j* ∈ [*N* ] \ {*i*} and *N* ∈ ℕ. The *X*_1_,…, *X*_*N*_ are exchangeable for each *N* ℕ; when *E*_1_ occurs the index *i* is picked uniformly at random from [*N* ]. The coalescents resulting from Definition 3.10 are described in Theorem B; § 6.2 contains a proof of Theorem B.

#### Theorem B

(Coalescents under Definition 3.10). *Suppose a diploid population evolves according to Definitions 2.3 and 3.10 and that Assumption 3.3 holds. Then* {*ξ*^*n,N*^ ⌊ *t/c*_*N*_ ⌋; *t* ≥ 0} *converges, in the sense of convergence of finite-dimensional distributions, to* {*ξ*^*n*^} ≡ {*ξ*^*n*^(*t*); *t* ≥ 0} *as specified in each case*.

1. *Suppose ζ*(*N* )*/N* → 0. *Then* {*ξ*^*n*^} *is the Kingman-coalescent*.
2. *Suppose* 0 < *α* ≤ 1 *and ζ*(*N* )*/N ≩* 0 *(recall* (22) *in Definition 3.1). Then* {*ξ*^*n*^} *is the* Ω*-δ*_0_*-Beta*(*γ*, 2 − *α, α*)*-coalescent defined in Definition 2.7 with transition rates as in* (28).

*In both cases* (31) *is in force* 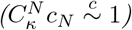 *with* 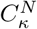 *as in* (23).

As in Corollary 3.9, restricting to completely dispersed states we have weak convergence on *D*([0, ∞), *ℰ*_2*n*_). The proof of Corollary 3.11 is identical to the one of [Birkner et al., 2018, Corollary 1.2].

#### Corollary 3.11

(Weak convergence of {*ξ*^*n,N*^} on *D*([0, ∞ ), ℰ_2*n*_)). *As in Corollary 3.9, let* 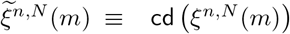 *for all m. Under the conditions of Theorem B* 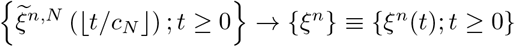 *weakly on D* ([0, ∞), ℰ_2*n*_) *as N* → ∞ *with* {*ξ*^*n*^} *as given in Theorem B*

### 3.3 Varying population size

Given the time-scaling (31), meaning that gene genealogies span on average numbers of generations proportional to the population size, it is plausible that the population size varies enough during the history of a sample to affect gene genealogies. Freund [2020] gives conditions for deriving time-changed Λ-coalescents describing gene genealogies in haploid panmictic populations with varying population size. We adapt the arguments for [Freund, 2020, Theorem 3] to diploid populations evolving according to Definition 2.3 and Definition 3.4 (or Definition 3.10).

Suppose 2*N*_*r*−1_ diploid potential offspring produced by *N*_*r*_ arbitrarily formed parent pairs survive to maturity in generation *r* where, for all *r* ∈ *t/c*_*N*_, *t* > 0 fixed, *N* ≡ *N*_0_, and with *c*_*N*_ being the coalescence probability for the fixed population size case,

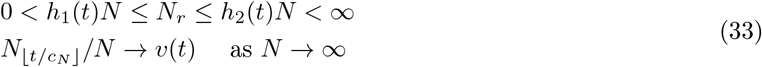

for some bounded positive functions *h*_1_, *h*_2_,*v* : [0, ∞) → (0, ∞) [Freund, 2020, Equation 4]. Since *c*_*N*_ is regularly varying (recall (31)) it holds that 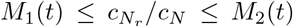 for all 0 ≤ *r* ≤ *t/c*_*N*_ for some *M*_1_(*t*), *M*_2_(*t*) ∈ (0, ∞) [Freund, 2020, Equation 13]. Moreover, writing 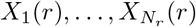 for the number of potential offspring produced in generation *r* (from whom 2*N*_*r*−1_ will be sampled to survive to maturity), it follows as in the proof of [Freund, 2020, Theorem 3] that (if (*x*_1_, *x*_2_,.. .) ∈ *o*_Σ_(*c*_*N*_ ) then 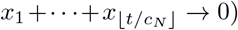

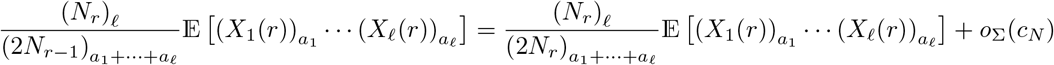

for *a*_1_ ≥ *a*_2_ ≥ … ≥ 1 [Freund, 2020, Equation 5]. Coupled with lim _*N*→∞_ *c*_*N*_ = 0, and Corollaries 3.9 and 3.11, it holds that 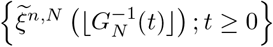 converges weakly to {*ξ*^*n*^} where 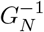 is as in [Freund, 2020, Equation 1]. Then, by (33) and (31) it follows that 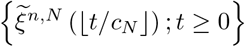 converges to {*ξ*^*n*^(*G*(*t*)); *t* ≥ 0}, where 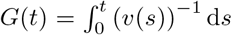 with *v* as in (33) [Freund, 2020, Lemma 4], and {*ξ*^*n*^ (*G*(*t*)) ; *t* ≥ 0} as given in Corollaries 3.9 resp. 3.11. Moreover, we have an analogue of [Freund, 2020, Lemma 1].

#### Lemma 3.12

(ℙ (*S*_*N,r*_ ≤ 2*N*_*r*−1_) vanishes). *Fix t* > 0. *Suppose* (*N*_*r*−1_ − *N*_*r*_) */N* → 0 *as N* → ∞ *for all* 0 ≤ *r* ≤ *t/c*_*N*_ . *Then there exists a constant* 0 < *c* < 1 *such that* ℙ 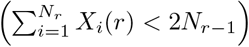 ≤ *c*^*N*^ *with N* = *N*_*0*_

*Proof of Lemma 3.12*. Write 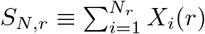. For 0 ≤ *s* ≤ 1 it holds that

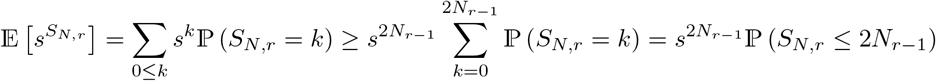

Write 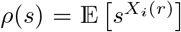. Then 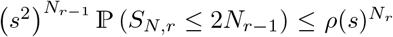 . Define *d*_*N,r*_ ≡ *N*_*r*−1_ − *N*_*r*_ and 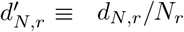 such that 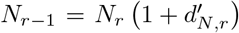. It follows that ℙ 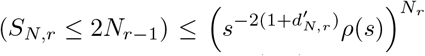 . Since *ρ*(1) = 1, and *ρ*^′^ (1) = E [*X*_*i*_(*r*)] > 2, there is an 0 < *s*_0_ < 1 and an *t:* > 0 such that 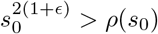.

Thus, the arguments for [Freund, 2020, Theorem 3] can be extended to Ξ-coalescents.

#### Theorem C

(Time-changed Ξ-coalescents). *Suppose a diploid population evolves as in Definition 2.3 and either Definition 3.4 or Definition 3.10. For any v* : [0, ∞ ) → (0, ∞) *there exist deterministically varying sizes* 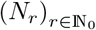 *such that* (33) *holds for the given v, and that* 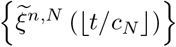 *converge weakly to* {*ξ*^*n*^ (*G*(*t*)) ; *t* ≥ 0} *where* 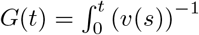 d*s. Here*, {*ξ*^*n*^ (*t*); *t* ≥ 0} *is as given in Theorem A or B*.

## 4 Comparing processes

Recall that {*ξ*^*n,N*^ } ≡ {*ξ*^*n,N*^ (*r*); *r* ∈ ℕ_0_} denotes an (annealed) ancestral process (tracking the random ancestral relations of sampled gene copies in a finite population) with time measured in generations, and {*ξ*^*n*^} ≡ {*ξ*^*n*^(*t*); *t* ≥ 0} denotes a (annealed) continuous-time coalescent. In this section we use simulations to investigate how well a given coalescent {*ξ*^*n*^} is approximated by the ancestral process in the domain of attraction to {*ξ*^*n*^}. We will also compare annealed and quenched (see below) gene genealogies (using simulations). In both cases we will compare empirical means of functionals of processes. First, we compare annealed processes {*ξ*^*n*^} and {*ξ*^*n,N*^}, with {*ξ*^*n,N*^} in the domain of attraction of the given {*ξ*^*n*^}.

### 4.1 Annealed ancestral processes and coalescents

We write |*A*| for the number of elements in a given finite set *A*. For *i* = 1, 2,…, 2*n* − 1 consider the functionals (*n* represents the number of sampled diploid individuals yielding 2*n* sampled gene copies)

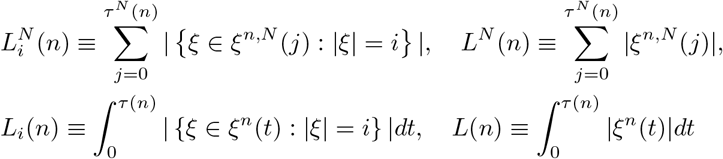

where *τ* ^*N*^ (*n*) ≡ inf {*j* ∈ ℕ : |*ξ*^*n,N*^ (*j*)| = 1}, *τ* (*n*) ≡ inf {*t* ≥ 0 : |*ξ*^*n*^(*t*)| = 1}, 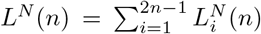, and 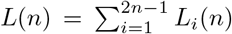 [Birkner et al., 2018, Section 1.1]. Interpreting {*ξ*^*n,N*^} and {*ξ*^*n*^} as trees one can interpret 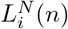 and *L*_*i*_(*n*) as the random length of branches supporting *i* leaves (Figure 1).

**Figure 1:**
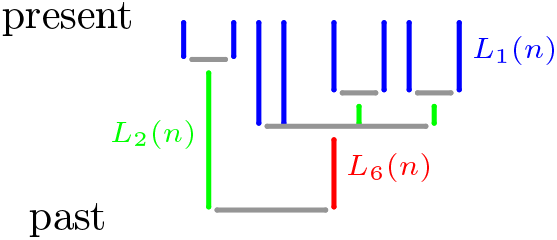
An example gene genealogy and the functionals *L*_*i*_(*n*); *L*_1_(*n*) is the sum of the length of the blue branches, and *L*_2_(*n*) the sum of the length of the green branches.

We assume that *ξ*^*n,N*^ (0) = {{1, 2},…, {2*n* − 1, 2*n*}}, i.e. we sam{ple *n*}diploid individuals and so 2*n* gene copies partitioned at time 0 as just described. When sampling {ξ^*n,N*^} we track the pairing of gene copies in diploid individuals; gene copies residing in the same diploid individual necessarily disperse (recall Definition 2.3 and Illustration 2.4).

Define, for *i* = 1, 2,…, 2*n* − 1,

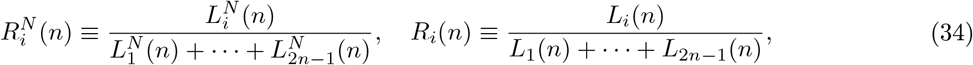

where *L*^*N*^ (*n*) ≥ 2*n* +2 and *L*(*n*) > 0 both almost surely. The functionals we will be concerned with are

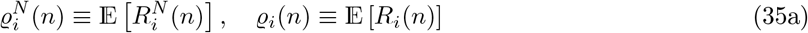

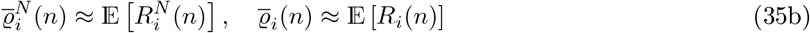

with 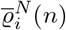 resp. 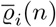 in (35b) denoting the corresponding empirical means computed from a finite number of independent realisations (using simulations) of 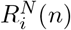 and *R*_*i*_(*n*), respectively.

When evaluating 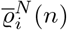 we keep track of the configuration of blocks in individuals (recall Definition 2.3 and Remark 2.4). Due to the instantaneous complete dispersion of blocks of {*ξ*^*n*^}, we assume *ξ*^*n*^(0) = {{1},…, {2*n*}} and *ξ*^*n*^(*t*) ∈ *ℰ*_2*n*_ for all *t* ≥ 0, i.e. the blocks of any partition *ξ* ∈ {*ξ*^*n*^} are always assumed to be completely dispersed. For any given coalescent {*ξ*^*n*^ } and an ancestral process {*ξ*^*n,N*^ (⌊*t/c*_*n*_⌋); *t* ≥ 0 in the domain-of-attraction of {*ξ*^*n*^} it should hold that 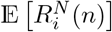 predicted by {*ξ*^*n,N*^ } converges to E[*R*_*i*_(*n*)] as predicted by{ *ξ*^*n*^ } . Comparing 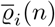 and 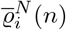 is then a way to see how well {*ξ*^*n,N*^} and {*ξ*^*n,N*^}agree.

For evaluating 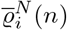 we will consider a population of constant size 2*N* evolving according to Definition 2.3 with the law for the random number *X* of potential offspring of an arbitrary parent pair being

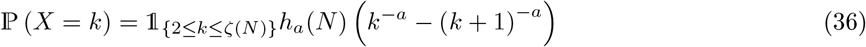

where *h*_*a*_(*N* ) is such that ℙ (*X* ∈ 2,…,*ζ*(*N* ) ) = 1. It then holds that *X*_1_ + … + *X*_*N*_ ≥ 2*N* almost surely.

Moreover, *h*_*a*_(*ζ*(*N* )) → 2^*a*^ as *ζ*(*N* ) → ∞ . The law in (34) is a special case of the one in (25).

Using the upper bound in (45) in Lemma 6.1 we see

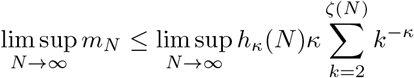

Similarly from the lower bound in (45) we see

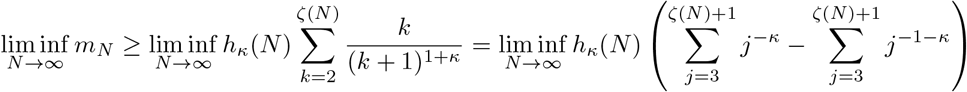

When *κ* = 2 it then holds that (recalling the 2-series 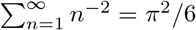)

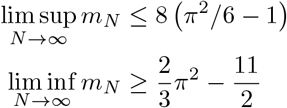

Recalling (36), and noting that 2*π*^2^*/*3 − 11*/*2 < 2, we approximate *m*_∞_ with 2 < m < 8 ((*π*^2^*/*6) −1) when *κ* = 2 (recall *X*_*i*_ ≥ 2 for all *i* ∈ [*N* ] by (36)).

When *κ* > 2 similar calculations show that

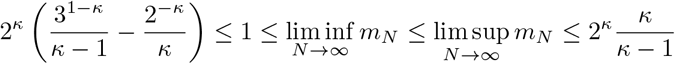

Then, recalling again (36) we approximate *m*_∞_, when *κ* > 2, with

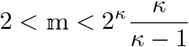

It follows from Case 2 of Theorem A, and Case 2 of Theorem B that the transition rate of a size-ordered (*k*_1_,…, *k*_*r*_)-merger for all *r* ∈ [4] is (recall (28) and (29a)–(29e))

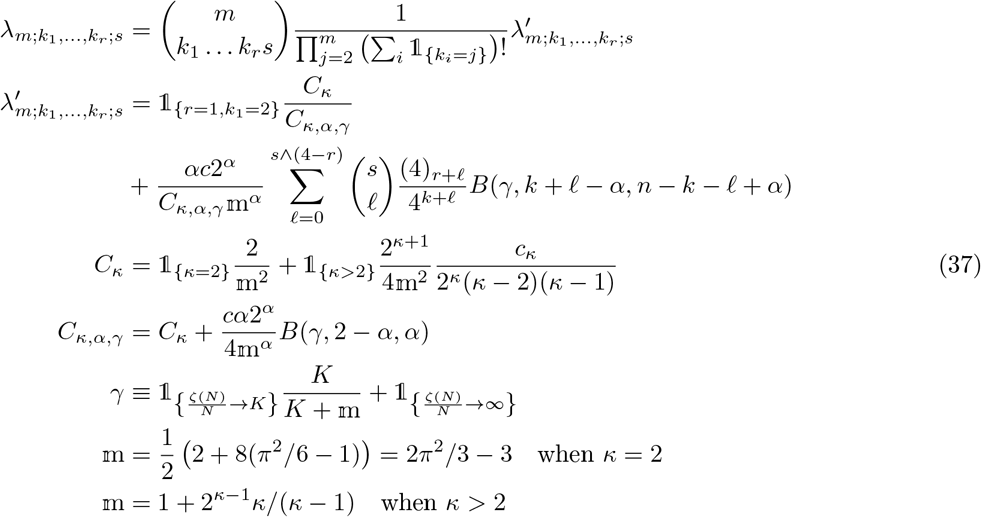

In (37) 0 < *α* < 2, 0 < *γ* ≤ 1, *c* > 0, *κ* ≥ 2.

Examples of 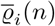 (recall (35b)) when {*ξ*^*n*^} is the Ω-*δ*_0_-Beta(*γ*, 2 −*α, α*)-coalescent (transition rates as in (37)) are in Figure 2. The Ω-*δ*_0_-Beta(*γ*, 2 −*α, α*)-coalescent can predict a range of different site-frequency spectra, including *U* -shaped spectra similar to that observed in the highly fecund diploid Atlantic cod [Árnason et al., 2023]. Morever, Figure 2 clearly shows the effect of the upper bound *ζ*(*N* ) (through *γ*; recall (29a)) on the site-frequency spectrum.

**Figure 2:**
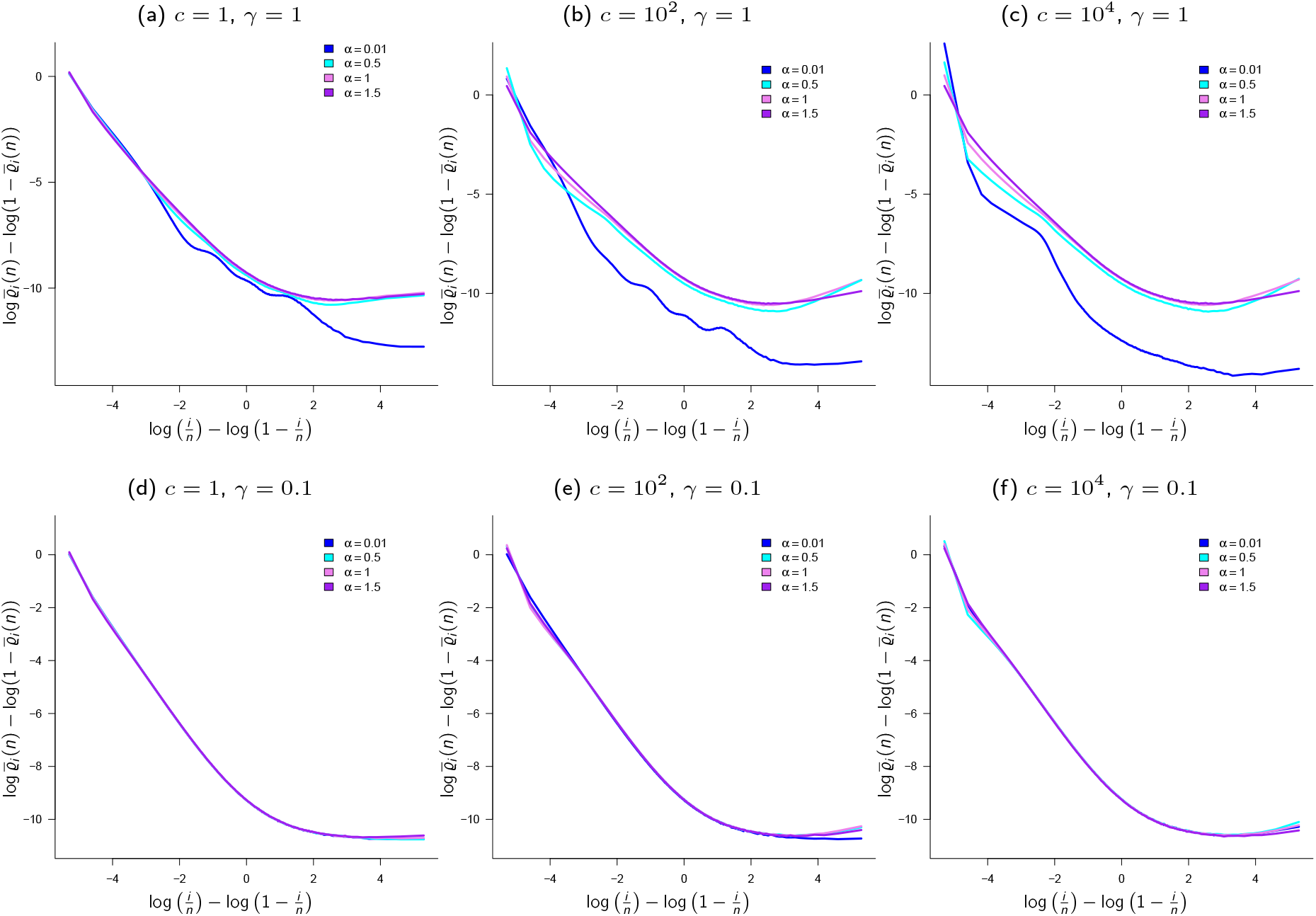
The Ω-*δ*_0_-Beta(*γ*, 2 −*α, α*)-coalescent. The empirical mean 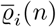 of *R*_*i*_(*n*) (recall (34), (35a), (35b)) when {*ξ*^*n*^} is the Ω-*δ*_0_-Beta(*γ*, 2 −*α, α*)-coalescent; when *n* = 100, *κ* = 2, *c, α*, and *γ* as shown; see Appendix B for a brief description of the algorithm for sampling from the Ω-*δ*_0_-Beta(*γ*, 2 −*α, α*) coalescent; results from 10^6^ experiments

However, there is discrepancy between 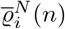 and 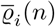 (recall (35a) and (35b)) when {*ξ*^*n*^} is the Ω-*δ*_0_-Beta(*γ*, 2 *−α, α*)-coalescent (Figure 3). In Figure 3 we compare 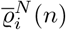 (recall (35b); blue and cyan lines) to 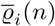 (violet and purple lines) when the population evolves as in Definition 2.3 and Definition 3.10 (Figure 3a) resp. Definition 3.4 (Figure 3b) with numbers of potential offspring distributed as in (36). One reason might be that {*ξ*^*n,N*^}converges slowly to {*ξ*^*n*^}, in particular for small . Should {*ξ*^*n*^} be a good approximation of {*ξ*^*n,N*^} only for population sizes exceeding realistic ones (the population size used in Figure 3 is likely nowhere near that), the usefulness of coalescent-based inference methods will then be left in doubt. In contrast, the agreement between Kingman and Wright-Fisher trees is in general quite good [Fu, 2006]. Figure 3 also reveals a qualitative difference within 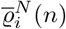 depending on the pre-limiting model (Definitions 3.10 and 3.4); the peaks in the graph of 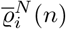 (blue lines in Figure 3a) are absent in Figure 3b. Similarly, the peaks in the graphs for 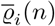 in Figure 3a are absent in Figure 3b.

**Figure 3:**
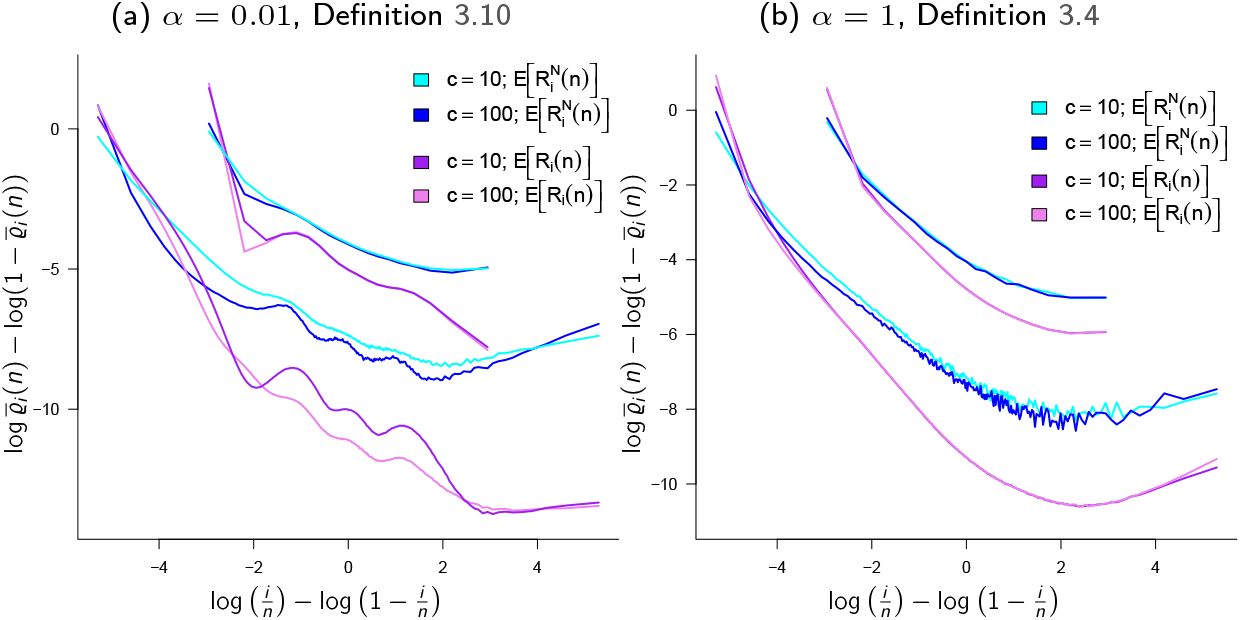
The Ω-*δ*_0_-Beta(*γ*, 2 −*α, α*)-coalescent. Comparing 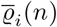 (recall (35a) and (35b)) and 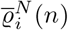 when {*ξ*^*n*^} is the Ω-*δ*_0_-Beta(*γ*, 2 *α, α*)-coalescent and when the population evolves according to Definitions 2.3 and 3.4 and 3.10 and (36) with *N* = 2500, *ζ*(*N* ) = *N* log *N, α* as shown, *κ* = 2, *ζ*(*N* ) = *N* log *N*, sample size *n* = 100, *n* = 10, *γ* = 1; *c* as shown; with *ε*_*N*_ as in (53) and 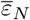 as in (61)

Recall from Case 3 of Theorem A that convergence to the Ω-*δ*_0_-Poisson-Dirichlet(*α*, 0)-coalescent as the limit of {*ξ*^*n,N*^} determined by Definitions 2.3 and 3.4, and (36), requires the rather strong (in the sense of being applicable to real populations; recall that 2*N* is the population size) assumption that *ζ*(*N* )*/N* ^1*/α*^ → ∞. Thus, we hesitate to claim that the Ω-*δ*_0_-Poisson-Dirichlet(*α*, 0)-coalescent is relevant for explaining population genetic data, even for broadcast spawners. Nevertheless, we record in Figures 6 and C1 examples of 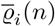 (recall (35a) and (35b)) for the Ω-*δ*_0_-Poisson-Dirichlet(*α*, 0)-coalescent.

### 4.2 Annealed and quenched ancestral processes

The annealed empirical means 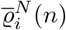 are obtained by (implicitly) averaging over the unknown ancestral relations of the sampled gene copies. An alternative way of estimating mean branch lengths is to condition on a given population ancestry (the ancestral relations of gene copies in the population). Given a population ancestry the gene genealogy of the gene copies in a given sample is complete and fixed, it is quenched. We let 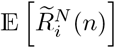 denote the quenched mean of relative branch lengths when averaging over population ancestries. Our approach is different from the one of Diamantidis et al. [2024], who consider the average over gene genealogies within one fixed population pedigree (the ancestral relations of diploid individuals).

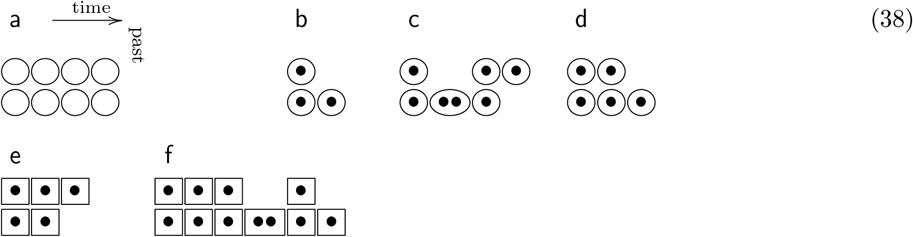

In (38b–d) are 3 possible gene genealogies of 2 gene copies (•) all going through the same pedigree (38a) (the ancestry of diploid individuals (○); from left to right is into the past). In contrast to restricting to a single pedigree, we average over population ancestries (a record of the ancestral relations of gene copies), recording the branch lengths of one complete sample gene genealogy per population ancestry (38e–f; the clusters of □ represent population ancestries); any sample of gene copies shares one gene genealogy within one and the same population ancestry. There are many possible gene genealogies within a fixed pedigree. Moreover, as Diamantidis et al. [2024] show, restricting to a fixed pedigree matters for what the limiting process (in a large population) will look like [Diamantidis et al., 2024, Theorem 1]. The view here is that the natural approach would be to condition on a population ancestry, read 1 complete sample tree from each given ancestry, and average the sample trees over independent ancestries. It remains to be seen what the limiting quenched coalescent then looks like, and if it is any different from the annealed coalescent.

We clarify the bookkeeping method for keeping track of the ancestral relations of the gene copies in the population. For each gene copy in the population we record the level it occupies, the time (generation) it occupies the given level, and the level occupied by the gene copy’s immediate ancestor (the value). Using a haploid panmictic population for an example, the offspring of the copy on level ℓ at time *g* all have value ℓ, while the gene copy itself derives from the one occupying level *i* at time *g* − 1 (see (39))

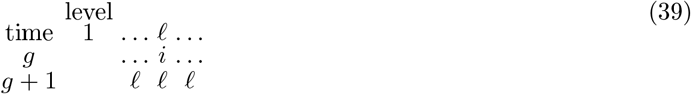

A diploid individual is seen as a pair of gene copies; for some *i* ∈ [2*N* ] gene copies occupying levels 2*i* − 1 and 2*i* at a given time form an individual. An example ancestry of a diploid panmictic population of 4*N* gene copies is shown in (40).

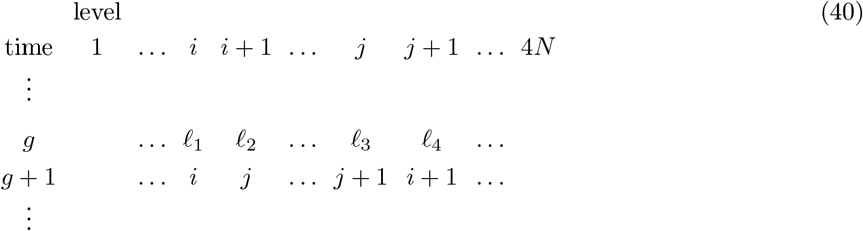

In (40) an individual whose gene copies occupy levels *i* and *i* + 1, and another individual whose gene copies occupy levels *j* and *j* +1 at time *g* form a parent pair; the gene copies of the surviving offspring of this parent pair ( {*i, i* + 1 }, {*j, j* + 1 }) will then have values taken from {*i, i* + 1, *j,j* + 1 }. We exclude selfing such that one gene copy of an offspring of {( *i, i* + 1 }, {*j, j* + 1 }) will sample its value from {*i, i* + 1 }. The other gene copy of the given offspring will sample its value from {*j, j* + 1 }. The parent gene copies themselves have values (immediate ancestors)_*l*1_,…, *l*_4_ as shown in (40); the gene copy on level *i* at time *g* derives from the one occupying level _1_ at time *g* − 1, and so on. In this way we record the population ancestry. Given a population ancestry the complete sample tree of a given sample of gene copies is fixed, as illustrated in (41).

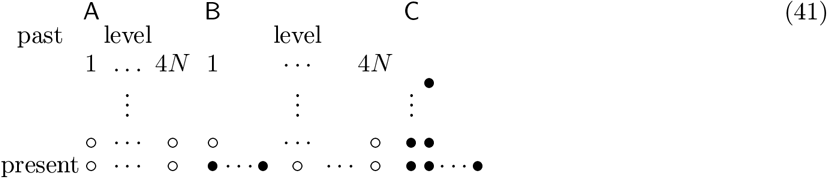

In (41) we broadly illustrate the process for generating a quenched gene genealogy. In (41A) we record a single population ancestry (the ancestral relations of the gene copies in the entire population) with denoting the gene copies (recall (40)); once the identity of the sampled gene copies (shown as in (41B)) is known all that remains is to track the fixed complete sample tree (41C). The empirical means 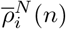 of quenched relative branch lengths are then obtained by repeating the method shown in (41), each time starting with a new population ancestry and reading 1 complete sample tree from each ancestry (see also Appendix E).

The annealed ancestral process is sampled by sampling offspring numbers and merging blocks assigned to the same parent chromosome (see Appendix D); the complete sample tree in the sense of the quenched method is never generated. We track block sizes of sampled blocks (and of blocks ancestral to the sampled blocks) and the partitioning of blocks in individuals as we transition backwards in time. The examples in (42) show 2 possible transitions for 4 blocks labelled *a, b, c*, and *d* all in separate diploid offspring of a given parent pair.

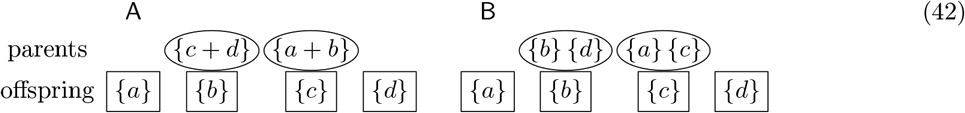

In (42A) blocks *c* and *d* merge, and blocks *a* and *b* merge in an example of a simultaneous merger; (42B) is an example of an instantaneous state in which blocks end up in the same diploid individual without merging. In (42A) we would continue with 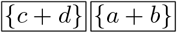, and with 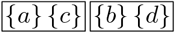 in (42B).

The means 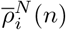 are evaluated for a diploid panmictic population of constant size 2*N* diploid individuals (4*N* gene copies) by evolving the population forward in time as in Definition 2.3 and (36) and recording the ancestry of the gene copies (recall (40)). Every now and then (meaning that we increment the population ancestry for a fixed number of generations in between drawing a sample) we randomly sample *n* diploid individuals (and so 2*n* gene copies), each time tracing the ancestry of the sampled gene copies, until a sample is obtained whose gene copies have a common ancestor (sampled gene copies without a common ancestor are discarded). The realised gene genealogy (gene tree) tracing the ancestry of the sampled gene copies is then said to be *complete*. The gene tree of the sampled gene copies is then fixed (for the given population ancestry and sample), and all that remains to do is to read the branch lengths off the fixed tree (recall (41)). Repeating this a given number of times, each time starting from scratch with a new population, gives us the empirical mean 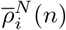, an estimate of 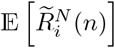 (see Appendix E).

In Figure 4 we compare 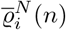 (an estimate of 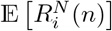 blue lines) and 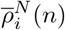 (an estimate of 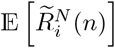 red lines) when the population evolves as in Definitions 2.3 and 3.4, and (36). When ζ(*N* ) = 2*N* (Figure 4a) then 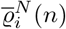 and 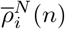 broadly agree; however when ζ(*N* ) = 4*N* ^2^ (Figure 4b) 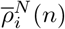 and 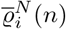 are qualitatively different (Figure E1 in Appendix E contains further examples).

**Figure 4:**
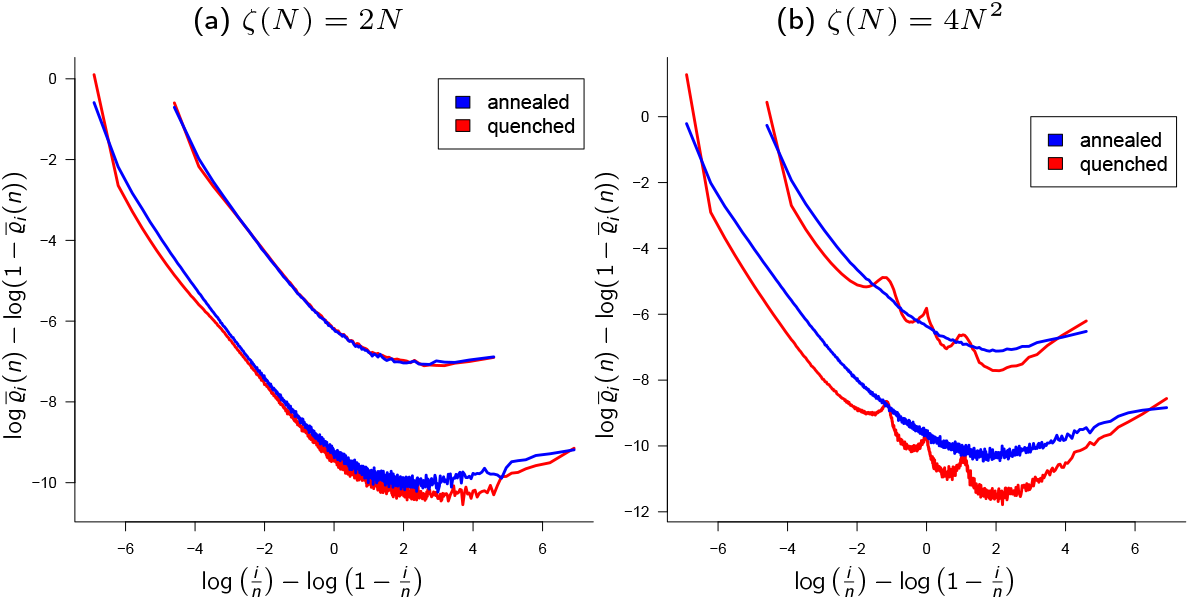
Annealed and quenched gene genealogies. Comparing 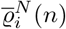 (an estimate of 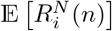 ; blue lines) and 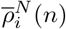 (an estimate of 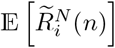 ; red lines) when the population evolves according to Definitions 2.3 and 3.4, and (36), for *N* = 250 (population size 2*N* of diploid individuals), upper bound ζ(*N* ) as shown, *ε*_*N*_ = 0.1, *α* = 1, *κ* = 2; sample size *n* = 50 and *n* = 500 diploid individuals; results from 10^5^ experiments. Appendices D and E contain brief descriptions of the sampling algorithms

### 4.3 Annealed time-varying coalescents

In Theorem C we extend [Freund, 2020, Theorem 3] to Ξ-coalescents where the time-change is independent of *α*. In Figure 5 we give examples of the effect of exponential population growth (time-change function *v*(*t*) = *e*^−*ρt*^) [Donnelly and Tavaré, 1995] when {*ξ*^*n*^ }is the Ω-*δ*_0_-Beta(*γ*, 2 −*α, α*)-coalescent, and in Figure 6 when {*ξ*^*n*^ } is the Ω-*δ*_0_-Poisson-Dirichlet(*α*, 0)-coalescent. Population growth extends the relative length of external branches of the examples of Ξ-coalescents studied here, similar to the effect on Kingman trees [Donnelly and Tavaré, 1995].

**Figure 5:**
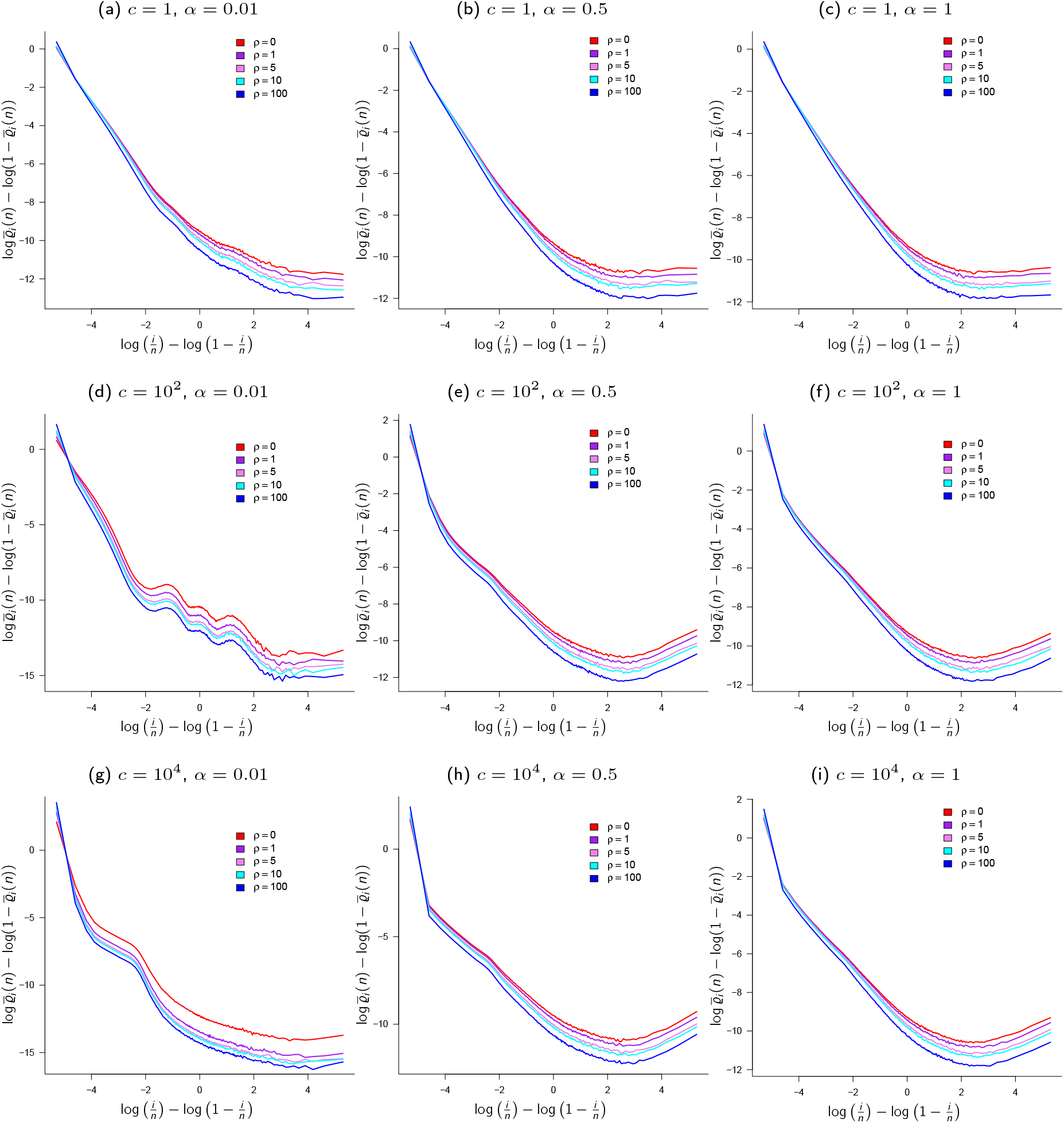
A (time-changed) Ω-*δ* -Beta(*γ*, 2 *α, α*)-coalescent. Empirical means 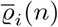 of E [*R*_*i*_ (*n*)] (recall (35a)) when {*ξ*^*n*^} is the time-changed Ω-*δ* -Beta(*γ*, 2 − *α, α*)-coalescent with time-change 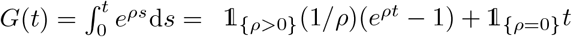; *γ =* 1,*α,c*,ρ as shown for *n* = 100; 10^5^ experiments

**Figure 6:**
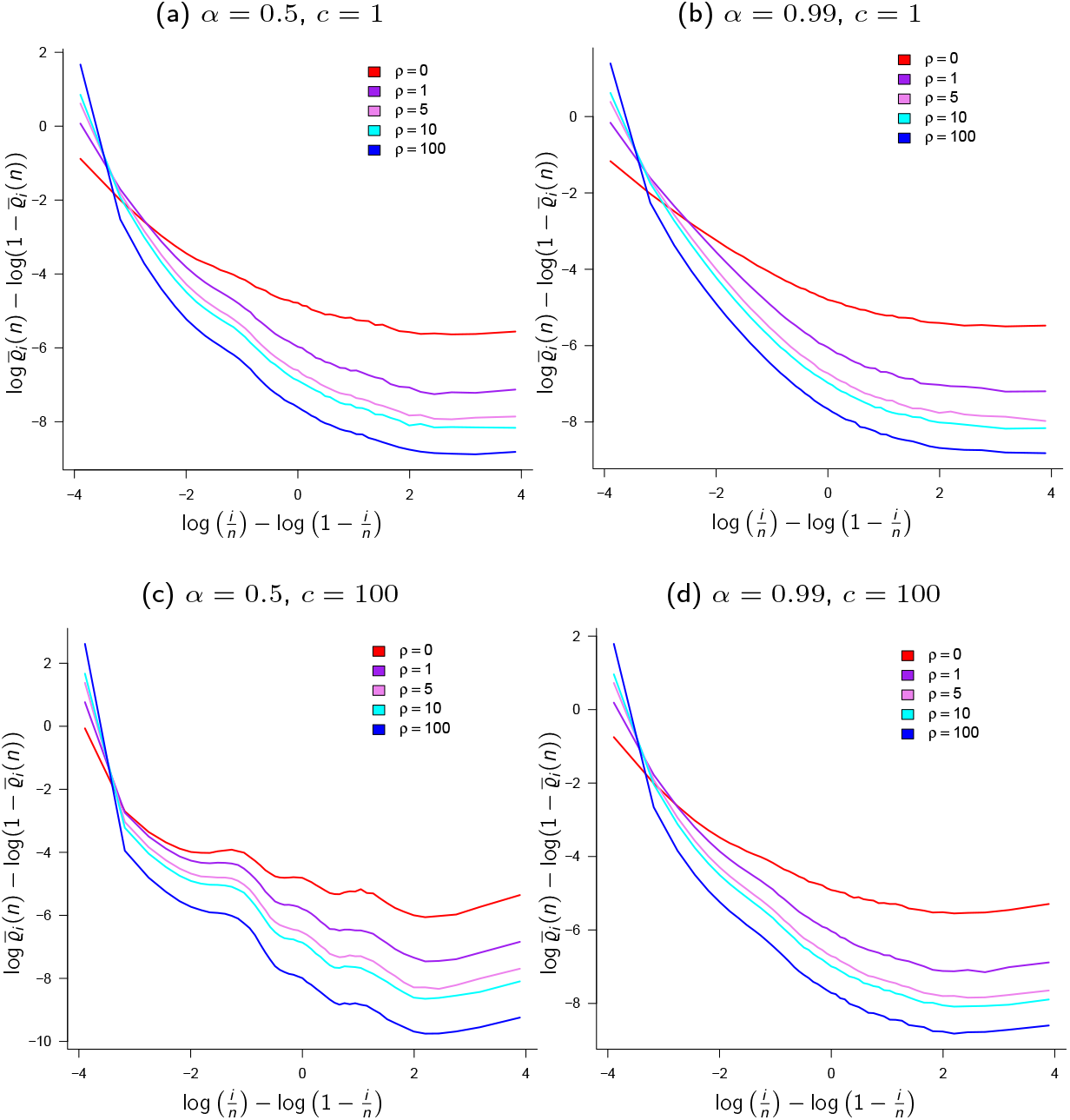
A (time-changed) Ω-*δ*_0_-Poisson-Dirichlet(*α*, 0)-coalescent. Empirical means 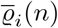 (recall (35a)) when {*ξ*^*n*^} is the time-changed Ω-*δ* -Poisson-Dirichlet(*α*, 0)-coalescent with time-change 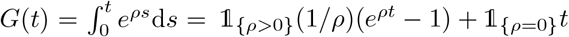 with *ρ, α, c* as shown, *κ* = 2; results from 10^5^ experiments

Since (31) is in force, such that gene genealogies (on average) span lengths of time proportional to *N* generation, it is plausible that some components of (25) may vary over time. In Figure 7 we give examples taking *γ*_*t*_ to be a simple step-function. Here, we take 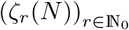 to be a sequence of cutoffs (with ζ_*r*_(*N* ) being the cutoff in (25) at generation *r* into the past) such that 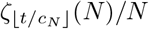 converges uniformly to some bounded positive function. Assuming a constant population size, our calculations for Case 2 of Theorem A, and Case 2 of Theorem B then show that the ancestral process {*ξ*^*n,N*^} converges (in finite-dimensional distributions) to a time-varying Ω-*δ*_0_-Beta(*γ*_*t*_, 2 − *α, α*)-coalescent.

**Figure 7:**
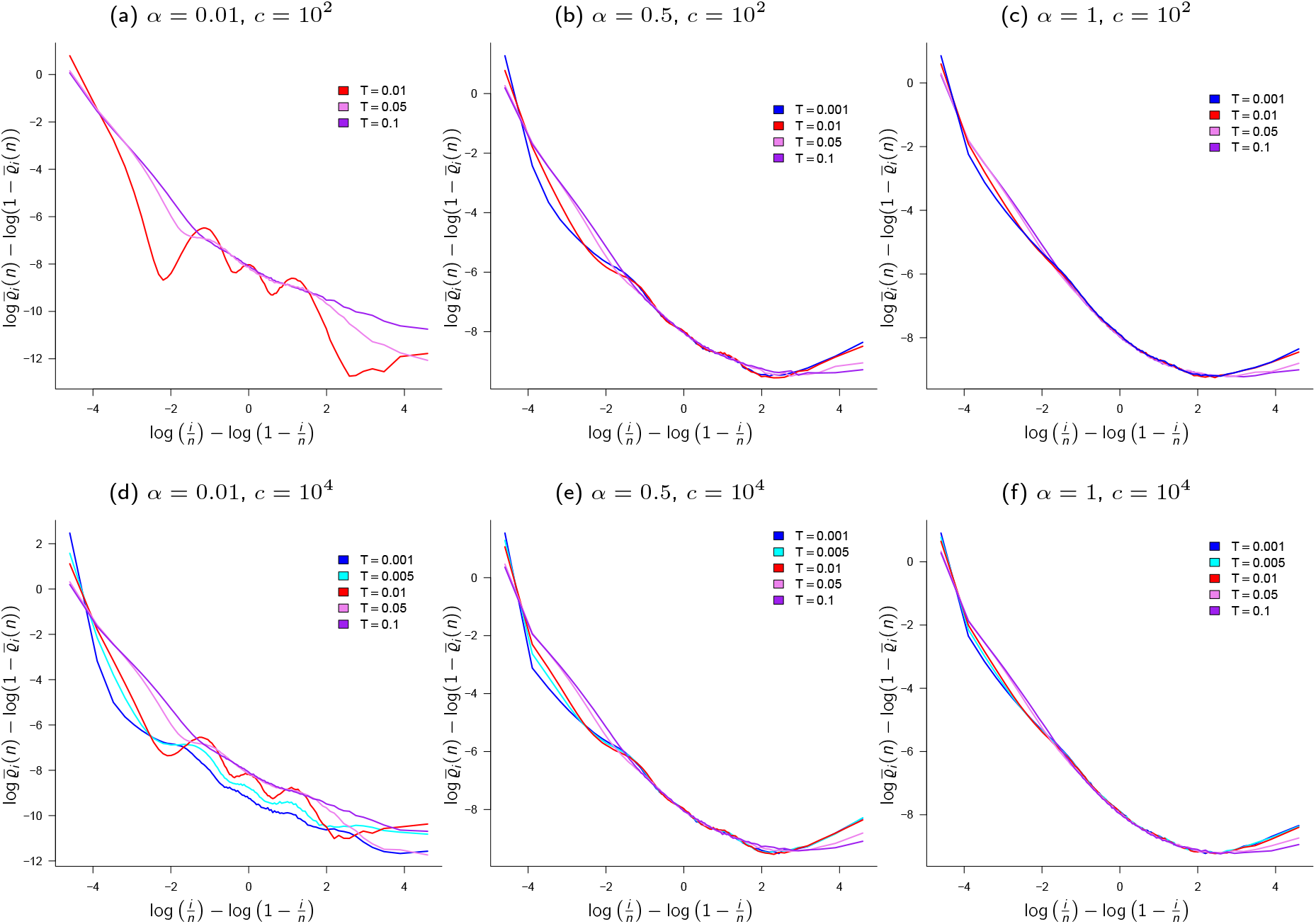
A time-changed Ω-*δ*_0_-Beta(*γ*_*t*_, 2 − *α, α*)-coalescent. Empirical means 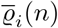 estimating E [*R*_*i*_(*n*)] (recall (35b)) when {*ξ*^*n*^} is the time-varying Ω-*δ*_0_-Beta(*γ*_*t*_, 2 − *α, α*)-coalescent with *γ*_*t*_ a step function *γ*_*t*_ = 1 _{0≤*t*≤*T* }_ 0.1 + 1 _{*t*>*T* }_ with *T* as shown, *κ* = 2; results from 10^5^ experiments

## 5 Conclusion

Our main results are *(i)* continuous-time coalescents that are either the Kingman coalescent or simultaneous multiple-merger coalescents as specific families of Beta- or Poisson-Dirichlet-coalescents including an atom at zero; *(ii)* in arbitrarily large populations time is measured in units proportional to either *N/* log *N* or *N* generations; *(iii)* it follows that population size changes (satisfying specific assumptions) lead to time-changed coalescents where the time-change is independent of the skewness parameter *α*; *(iv)* in scenarios with increased effect of sweepstakes (e.g. ζ(*N* )*/N* → ∞) the annealed and quenched empirical means 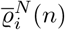 and 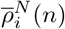 are quite different.

Based on our construction of a population model where the population evolves as in Definition 2.3 and incorporates sweepstakes as in Definitions 3.4 or 3.10 with an upper bound on the number of potential offspring (recall (25)) the resulting coalescent is driven by a measure of the form (recall (22) in Definition 3.1)

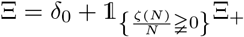

where Ξ_+_ is a finite measure on Δ_+_ (recall (9)). In our formulation ζ(*N* ) determines if the limiting coalescent admits multiple mergers or not. For comparison, when the *X*_1_,…, *X*_*N*_ are allowed to be arbitrarily large and with a regularly varying tail (recall (8)) as in [Birkner et al., 2018, Equation 26] then Ξ = 1_{*α*≥2}_*δ*_0_ + 1 _1<*α*<2}_Ξ_+_ [Birkner et al., 2018, Proposition 2.5]. When Ξ_+_(Δ_+_) < 1 one can assign the mass 1 − Ξ_+_ (Δ_+)_ to (0,.. .) [Birkner et al., 2018, Section 1]. We elect to explicitly model scenarios as in Definitions 3.4 and 3.10 where most of the time small families are generated (corresponding to most of the time sampling the zero atom (0,.. .)), but occasionally there is an increased chance of large families (corresponding to sampling elements from Δ_+_). Our formulation enables us to compare {*ξ*^*n,N*^} and {*ξ*^*n*^}, and the annealed and quenched processes, using the empirical means 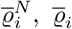, and 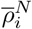. In addition, we show that explicitly incorporating a random environment in the sense of Definitions 3.4 and 3.10 alters the timescaling (recall (31) and (23)). The unit of time in these models has relevance for explaining observed genetic variation; one should be able to recover the mutations used to obtain parameter estimates for a given model without unrealistic assumptions about the population size or the mutation rate.

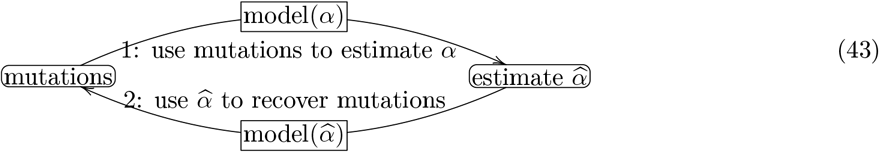

In (43) “model” refers to coalescent transition rates, the time-scaling (1*/c*_*N*_ ) used for passing to the coalescent limit, and assumptions about the mutation mechanism. Mutation rates (per site per generation) can be estimated using molecular methods [Milholland et al., 2017, Xue et al., 2009, Wooldridge et al., 2025, Seplyarskiy et al., 2023, Wang et al., 2026].

The models considered here (recall Definitions 2.3, 3.4, and 3.10) can be a basis for constructing ancestral influence graphs for diploid highly fecund populations [Koskela and Berenguer, 2019], and so for investigating the effects of elements such as population structure, range expansion, recurrent bottlenecks, and natural selection on genetic variation. Moreover, the time-scaling (31) implies, with gene genealogies now spanning (on average) time intervals proportional to (at least) *N/* log *N* generations, that some components of (25) may vary over time. Our numerical results should encourage comparison of processes as in § 4, and mathematical studies of quenched (conditional) gene genealogies.

## 6 Proofs

In this section we give proofs of Theorems A and B. First, we record a useful approximation of the bounds in (25).

### Lemma 6.1

(Eldon [2026], bounds on *k*^−*a*^ − (1 + *k*)^−*a*^). *Suppose* 0 < *a* ≤ 1 *and k* ∈ ℕ. *Then*

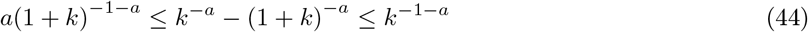

*If a* ≥ 1, *and k* ≥ 2, *it holds that*

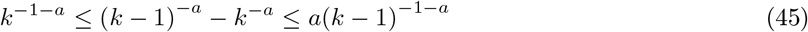

The following lemma is a straightforward extension of [Schweinsberg, 2003, Lemma 6; Equation 16]; recall that, if the population size is constant at 2*N* diploid individuals, it holds *ν*_1_ + · + *ν*_*N*_ = 2*N* .

### Lemma 6.2

(Relation between transition probabilities). *With ν*_1_,…, *ν*_*N*_ *and X*_1_,…, *X*_*N*_ *as in Definition 3.1 we have, with k*_1_,…, *k*_*r*_ ≥ 2 *and k* = *k*_1_ + · + *k*_*r*_ *for any r* ∈ ℕ *(recall S*_*N*_ *from* (20a))

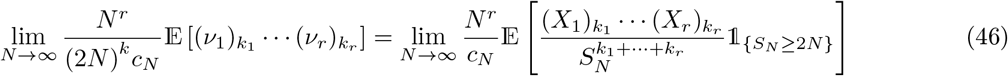

*in the sense that when one of the limits exists so does the other, and when they do exist they are equal. Moreover, as N* → ∞, *recalling c*_*N*_ *from Definition 2.2*,

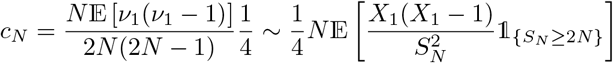

### Remark 6.3.

*The existence of* 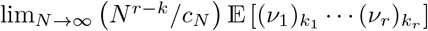 *implies the existence of*

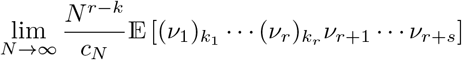

*for any s* ∈ ℕ *[Möhle and Sagitov, 2001, Lemma 3.5]*.

### Proposition 6.4

(Chetwynd-Diggle and Eldon [2026] approximating Σ_*k*_ *H*(*k*)(*G*(*k*) − *G*(*k* + 1))). *Suppose G and H are positive functions on* [1, ∞), *H monotone increasing, G monotone decreasing, and exists. For, m* ∈ ℕ, ≤ *m*,

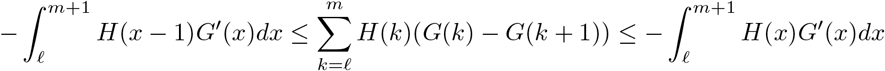

### Proposition 6.5

(Bounding Σ _*k*_ *k*(*k* − 1)(*k* + *M* )^−2^ (*k*^−*a*^ − (*k* + 1)^−*a*^)). *Write* 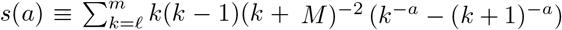 . *Suppose M* > 1 *is fixed*.

1. *When* 0 < *a* < 1 *or* 1 < *a* < 2 *it holds that*

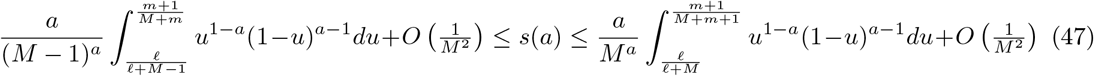
2. *When a* = 1 *we have*

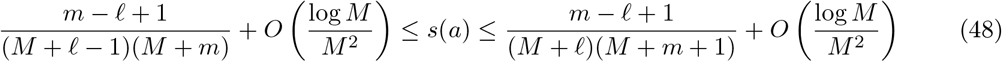
3. *When a* = 2 *we have*

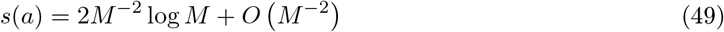
4. *When a* > 2 *it holds that*

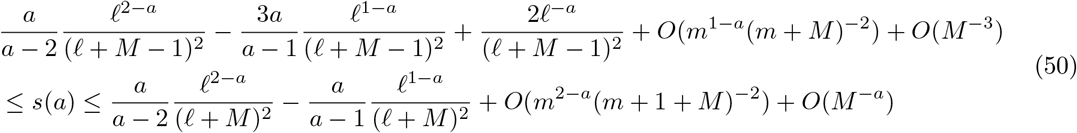

*Proof of Proposition 6.5*. Using Proposition 6.4 with *G*(*x*) = *x*^−*a*^, and *H*(*x*) = *x*(*x* − 1)*/*(*x* + *M* )^2^, we see

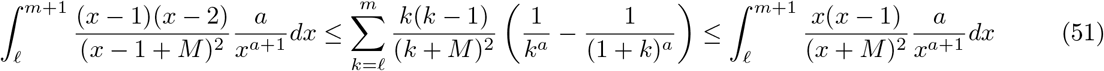

Evaluating the integrals in (51) using standard integral calculus techniques finishes the proof.

### Lemma 6.6

(P (*X*_1_ + · + *X*_*N*_ < 2*N* ) vanishes). *Suppose X*_1_,…, *X*_*N*_ *are independent non-negative integer-valued random variables such that* 2 < E [*X*_*i*_] < ∞ *for all i* ∈ [*N* ]. *A positive constant c* < 1 *then exists such that* P (*X*_1_ + · + *X*_*N*_ < 2*N* ) ≤ *c*^*N*^ *for all N* ∈ ℕ.

*Proof of Lemma 6.6*. The proof follows the on e of [Schweinsberg, 2003, Lemma 5]. Write *S*_*N*_ ≡*X*_1_+·· ·+*X*_*N*_ .

Let *ρ* : [0, 1] → [0, 1] be given by 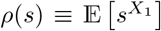 so that *ρ*^’^(1) = E [*X*_1_] > 2 and *ρ*(1) = 1. It then holds that *ρ*(*r*) < *r*^2^ for some number *r* ∈ (0, 1). Then 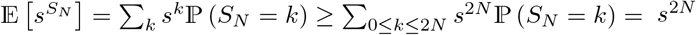 ℙ (*S*_*N*_ ≤ 2*N* ) so that P (*S*_*N*_ ≤ 2*N* ) ≤ (*s*^−2^*ρ*(*s*))^*N*^ for all 0 < *s* ≤ 1. Take *c* = *ρ*(*r*)*/r*^2^.

### 6.1 Proof of Theorem A

In this section we prove Theorem A. Recall Definition 3.4, and the notation in Definitions 2.1 and 3.1. We first give conditions on *ε*_*N*_ for *m*_∞_ from (21) to be finite.

#### Lemma 6.7

(Finite *m*_∞_). *Under the conditions of Theorem A with*

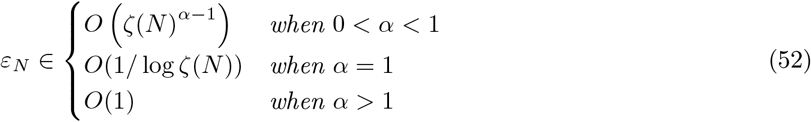

*as N* → ∞ *it holds that* lim sup_*N* →∞_ E [*X*_1_] < ∞.

*Proof of Lemma 6.7*. When 0 < *α* < 1 the upper bound in (44) gives, recalling 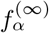 from (26),

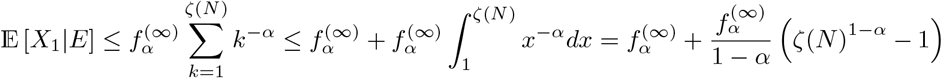

The boundedness of E [*X*_1_ |*E*^c^] follows by similar arguments. By Definition 3.4 E [*X*_1_] = E [*X*_1_ *E*] *ε*_*N*_ + E [*X*_1_| *E*^c^] (1 − *ε*_*N*_ ), hence choosing *ε*_*N*_ as in the lemma the result follows. The other cases in (52) are obtained using similar calculations.

#### Lemma 6.8

(*S*_*N*_ */*(*Nm*_*N*_ ) → 1 almost surely). *Suppose X*_1_,…, *X*_*N*_ *for each N* ∈ ℕ *are i.i.d. and distributed as in Definition 3.4 with ε*_*N*_ *as in Lemma 6.7 such that* lim sup_*N* →∞_ *m*_*N*_ < ∞ *for each N* ∈ ℕ *(then m*_*N*_ *converges to m*_∞_ ∈ (2, ∞) *by* (21) *and Remark 3.3). It holds S*_*N*_ */*(*Nm*_*N*_ ) → 1 *almost surely*.

*Proof of Lemma 6.8*. Recall *m*_*N*_ from] (21), *S*_*N*_ from (20a). Write 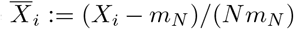 for all *i* ∈ [*N* ]; for *i* > *N* choose 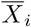 such 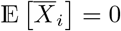, the 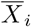 be independent, and 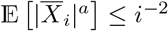 for some 1 < *a* < 2.

It follows that 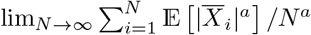 is finite, all the 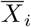 are independent, and so 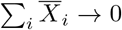 almost surely [Athreya and Lahiri, 2006, Theorem 8.4.6]. Since 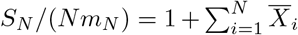, the lemma follows.

#### Remark 6.9

(Behaviour of *ζ*(*N* )). *Lemma 6.8 holds with m*_*N*_ *finite for each N. However, we are interested in cases where m*_∞_ *is finite. Therefore, we restrict the lemma to cases where* lim sup_*N*_ *m*_*N*_ < ∞. *For m*_∞_ *to be finite we require ζ*(*N* ) *not increase too quickly with N. Whenever ζ*(*N* )*/N* → ∞ *in Case 2 of Theorem A with* 1 ≤ *α* < 2 *we expect ζ*(*N* ) *to be at most of the form N*^*x*^ *for some fixed x* ≥ 1. *For example, α* = 1 *requires that x* < 2*a/*(*a* − 1), 1 < *a* < 2, *for Lemma 6.8 to hold*.

We verify (31) when 1 ≤ *α* < 2 under the conditions of Theorem A.

#### Lemma 6.10

(Verifying (31) when 1 ≤ *α* < 2). *Suppose the conditions of Theorem A hold, and that* 1 ≤ *α* < 2. *L<u>et</u> L* ≡*L*(*N* ) *be a positive function of N with L/N* → 0 *as N* → ∞. *Suppose* 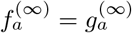, *and that* 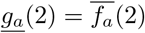 *(recall* (26)*). Take, for some c* > 0 *fixed (recall* 1_{·}_ *from* (5), *S*_*N*_ *from* (20a))

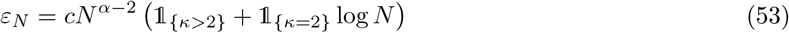

*Then* (31) *is in force, i.e*. 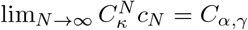 *with C*_*α,γ*_ *as in* (29c).

*Proof of Lemma 6.10*. Fix 0 < *δ*< 1 such that (1 − *δ*)*m*_*N*_ > 2 (recall (21) and Remark 3.3). Define, for each *N* ∈ ℕ,

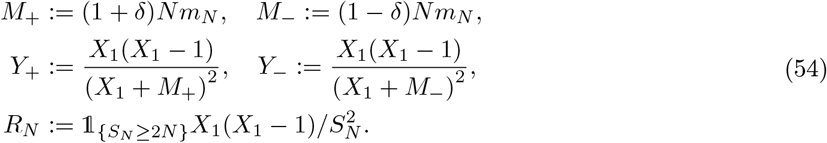

Fix *t:* > 0. Choosing *ε*_*N*_ (recall Definition 3.4) as in (52) gives *m*_∞_ < ∞ by Lemma 6.7. Using Lemma 6.8 we can adapt the arguments in the proof of [Schweinsberg, 2003, Lemma 13] to obtain

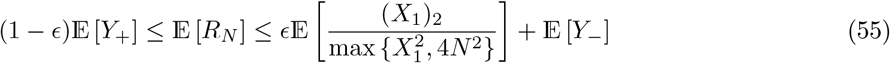

We will use Proposition 6.4 with *H*(*x*) = *x*(*x* − 1)(*x* + *M* )^−2^ and *G*(*x*) = *x*^−*a*^ with *M* either *M*_+_ or *M*_−_from (54) and *a* as given each time to approximate E [*Y*_−_] and E [*Y*_+_]. First, we check that

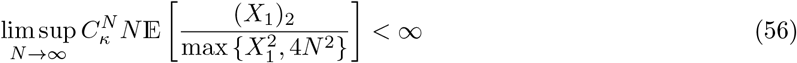

with *α* ≥ 1. Since (*X*_1_ + 2*N* )^2^ ≤ 4 max 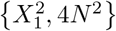 we have

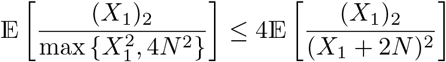

Using the upper bound in (45) in Lemma 6.1 we see, recalling event *E* from Definition 3.4,

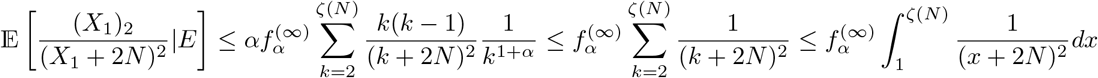

when *α* = 1; (56) follows with *ε*_*N*_ as in (53).

We approximate E [*Y*_−_] with *Y*_−_ as in (54). We see, using Proposition 6.4 with *G*^′^(*x*) = −*αx*^−*α*−1^, and *H*(*x*) = *x*(*x* − 1)(*x* + *M*_−_)^−2^,

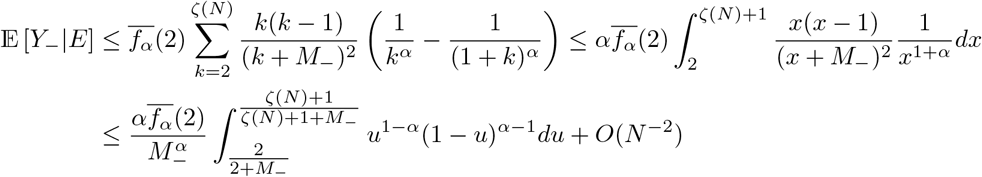

using the substitution *y* = *M* _−_*/*(*x* + *M*_−_) (so that *x* = *M*_−_*y*^−1^ − *M*_−_ and *dx* = −*M*_−_*y*^−2^*dy*) on *x*^1−*α*^(*x* + *M*_−_)^−2^*dx* and noting that (*x* + *M*_−_)^−2^*x*^−2^*dx* ≤ (*x* + *M*_−_)^−2^*x*^−*α*^*dx* over [1, ∞) when 1 ≤ *α* < 2. Write (recall *K* > 0 is fixed)

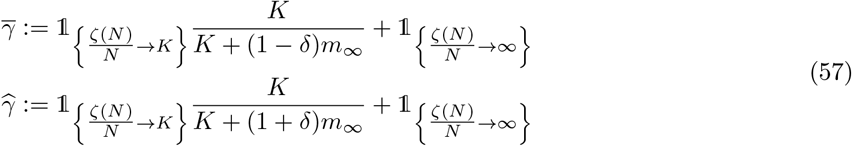

Using the assumption *L/N* → 0 with *ε*_*N*_ as in (53) and 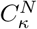 as in (23) we obtain

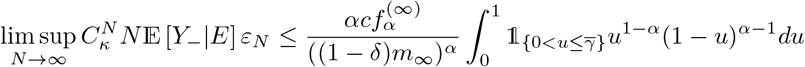

Analogous calculations give (recall 1 ≤ *α* < 2 by assumption)

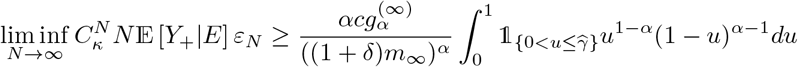

When *κ* = 2 we see, using Proposition 6.4 with *H*(*x*) = *x*(*x* − 1)(*x* + *M*_−_)^−2^ and *G*(*x*) = *x*^−2^,

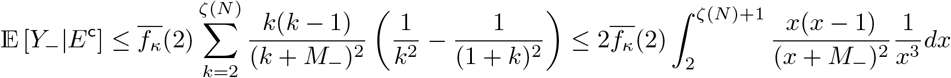

Integration by partial fractions gives

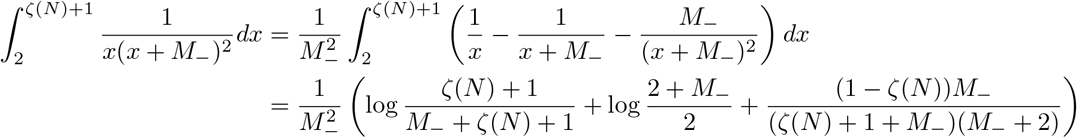

and we conclude, recalling 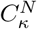 from (23) and that *κ* = 2,

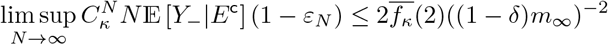

after checking that lim 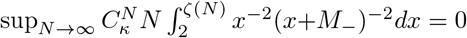 using integration by parts (where 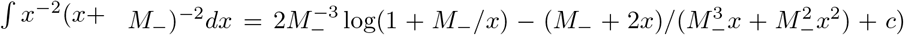 . Again using Proposition 6.4 with *H*(*x*) = *x*(*x* − _+_ ^−2^ and *G*(*x*) = *x*^−2^ we see

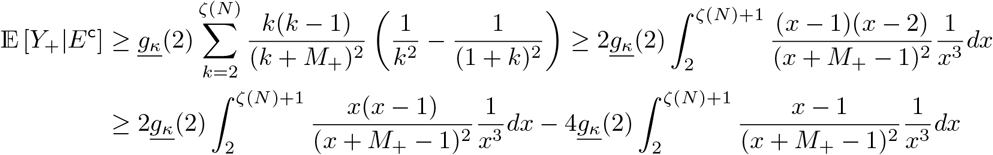

and we can conclude that, when *κ* = 2,

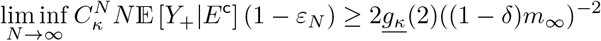

When *κ* > 2 we see, using Proposition 6.4,

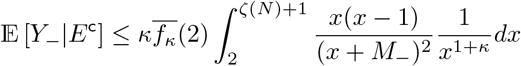

Suppose 2 < *κ* < 3. Integration by parts and the substitution *y* = *M*_−_*/*(*x* + *M*_−_) then give

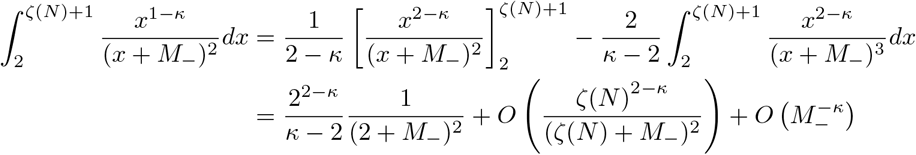

Iterating the calculation for *∫ x*^−*κ*^(*x* + *M*_−_)^−2^*dx* we conclude

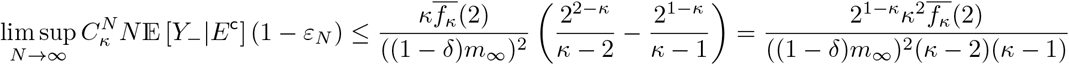

Similarly, when 2 < *κ* < 3, again using Prop 6.4,

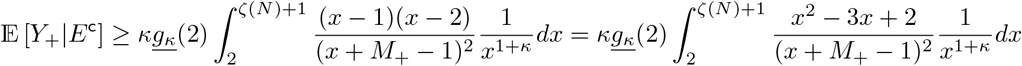

and we obtain

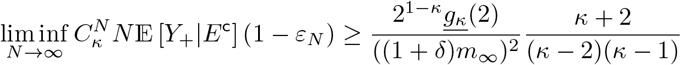

One checks in the same way that the same leading terms hold for *κ* ≥ 3. The lemma now follows from (7) and Lemma 6.2 after taking *ϵ* and *δ*to 0.

#### Lemma 6.11

(Convergence to the Kingman coalescent). *Under the conditions of Case 1 of Theorem A it holds that*

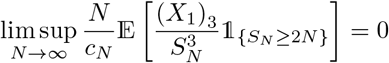

*Proof of Lemma 6.11*. On *S*_*N*_ ≥ 2*N* we have 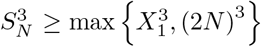 so that, on *E* (recall Definition 3.4) and with 0 < *α* < 1, using the upper bound in (44) in Lemma 6.1, and that |*x* + *y*|^*p*^ ≤ 2^*p*−1^ (|*x*|^*p*^ + |*y*|^*p*^) for reals *x, y* and *p* ≥ 1 [Athreya and Lahiri, 2006, Proposition 3.1.10*(iii)*, Equation 1.12]

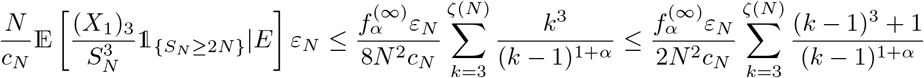

and the result follows when 0 < *α* < 1 assuming lim sup*N*_→∞_ *ε*_*N*_ */c*_*N*_ < ∞ as *N*→ ∞. The result when 1 ≤ *α* < 2 follows from a similar calculation. Convergence to the Kingman coalescent follows from Lemma 6.2 and [Schweinsberg, 2003, Proposition 2] (see [Möhle, 2000, Section 4]).

We check that the probability of two or more large families vanishes in a large population when 1 ≤ *α* < 2. However, as we will see, diploidy ensures that we retain the possibility of (up to 4-fold) simultaneous mergers.

#### Lemma 6.12

(The probability of two or more large families vanishes when 1 ≤ *α* < 2). *Under the conditions of Case 2 of Theorem A it holds that*

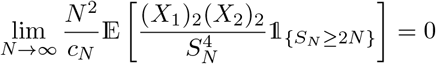

*Proof of Lemma 6.12*. We see, recalling that the *X*_1_,…, *X*_*N*_ for each *N* ∈ ℕ are i.i.d.,

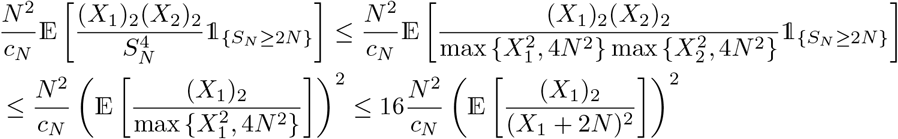

Recalling event *E* from Definition 3.4 we see, when 1 ≤ *α* < 2,

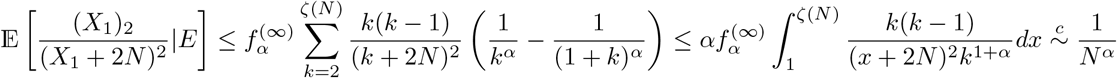

as *N* → ∞ (recall *ζ*(*N* )*/N* ≩ 0), using (45) in Lemma 6.1 and the substitution *y* = 2*N/*(*x* + 2*N* ) when 1 < *α* < 2. The result now follows from Lemma 6.10 with *ε*_*N*_ as in (53).

#### Lemma 6.13

(Identifying the measure Ξ_+_). *Under the conditions of Case 2 of Theorem A we have*

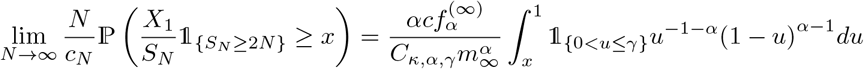

*for* 0 < *x* < 1 *and with C*_*κ,α,γ*_ *as in* (29c) *and γ as in* (29a) *and c from* (53) *and f*_∞_ *as in* (26).

*Proof of Lemma 6.13*. Let *ϵ, δ* > 0 be fixed with (1 − *δ*)*m*_*N*_ > 2 for all *N* . Then, by Lemma 6.8 we have for all *N* large enough that, with 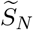 as in (20b) in Definition 3.1,

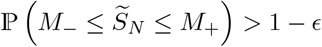

is in force where *M*_−_ and *M*_+_ are as in (54). Then, for any 0 < *x* < 1,

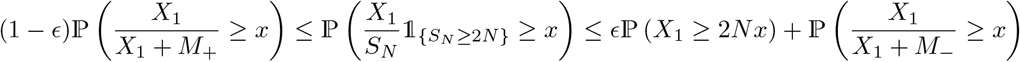

using the arguments for [Schweinsberg, 2003, Equations 44 and 45 in Lemma 14]. Recalling event *E* from Definition 3.4 we see

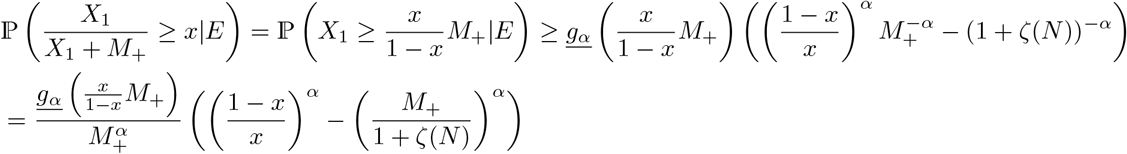

For *a* > 0 and 0 < *x* ≤ 1*/*(1 + *c*) for any constant *c* ≥ 0 we have (cf. [Chetwynd-Diggle and Eldon, 2026])

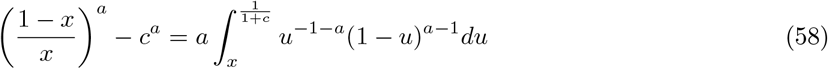

seen using the substitution *y* = (1 − *u*)*/u*. Write 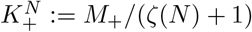 so that

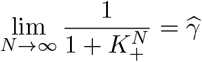

where 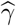 is as in (57). By Lemma 6.10 with *ε*_*N*_as in (53) we have (recall *C*_*κ,α,γ*_ from (29c))

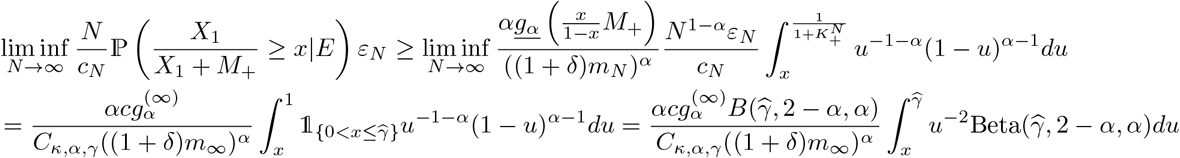

where Beta(*p*, 2 − *α, α*) is the law on (0, 1] with density as in (15). Analogous calculations give

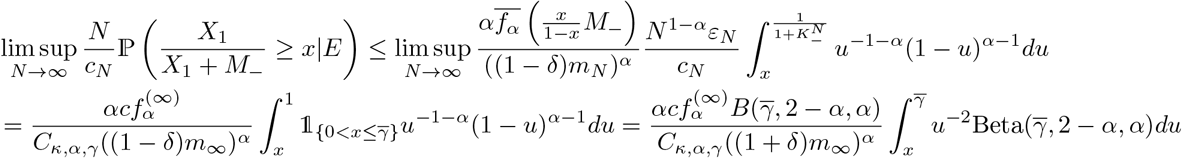

where 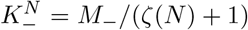 so that 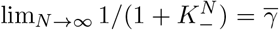 with 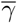 as in (57). The lemma follows after taking *ϵ* and *δ* to 0.

*Proof of Theorem A*. Case 1 follows from Lemmas 6.11 and 6.2, and from [Möhle and Sagitov, 2003, Theorem 5.4; Equation 2]. The proof of Case 2 follows as in the proof of [Birkner et al., 2018, Theorem 1.1] (in turn based on the results of Möhle [1998]); we only note that (recall *A* is the number of elements in a given finite set *A*)

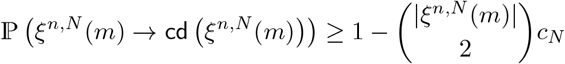

where *c*_*N*_ is the coalescence probability as in Defition 2.2 (recall (7)). Since *c*_*N*_ → 0 by Lemmas 6.10 and (46), and |*ξ*^*n,N*^ (*m*)| ≤ 2*n* < ∞, so that ℙ( *ξ*^*n,N*^ (*m*))→ cd (*ξ*^*n,N*^ (*m*))) → 1 as *N* → ∞, complete dispersion of ancestral blocks paired in the same diploid individual occurs instantaneously in the limit, and the limiting ε_2*n*_-process is valued (the set of partitions of [2*n*]).

By Lemma 6.10 and Lemma 6.2

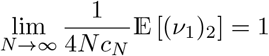

and the limits in (19a) are zero for all *r* ∈ ℕ \ {1} 1 by Lemma 6.12, the limits in (19a) hold for all *j* ∈ ℕ. Fix 0 < *x* < 1, and 0 < *ϵ* < *x*. Then, by the arguments for [Schweinsberg, 2003, Theorem 4c],

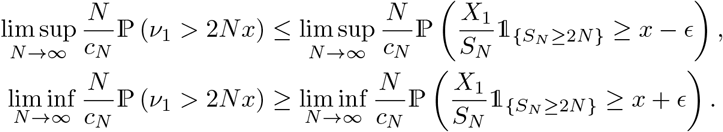

Then, after taking *ϵ* → 0, it holds 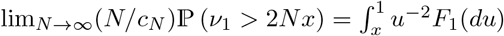, where *F*_1_ is the Beta(*γ*, 2 − *α, α*)-measure by Lemma 6.13. Furthermore,

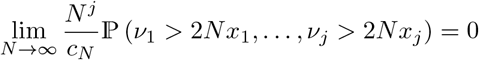

for all *j* ∈ {2, 3,.. .} by Lemma 6.12 and [Möhle and Sagitov, 2001, Equation18]. Hence, the limits in (19b) hold for all points of continuity for the measures *F*_*r*_ for all *r* ∈ ℕ. Thus, {*ξ*^*n,N*^ ( *t/c*_*N*_ ); *t* ≥ 0}→ {*ξ*^*n*^(*t*); *t* ≥ 0} in the sense of convergence of finite dimensional distributions.

We turn to Case 3 of Theorem A. The proof follows the one of [Schweinsberg, 2003, Theorem 4d] (see 774 also the proof of [Eldon, 2026, Theorem 3.5, Case 3]). Suppose 0 < *α* < 1, and *ζ(N)/N*^*1/α*^ → ∞. Let *Y*_*(1)*_ ≥ *Y*_(2)_ ≥ … ≥ *Y*_(*N*)_ be the ranked values of 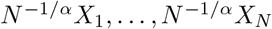 Write 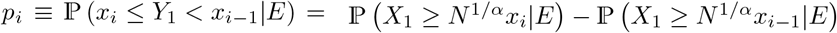 Using (27) we have, for 1 ≤ *i* ≤ *j*, recalling (26),

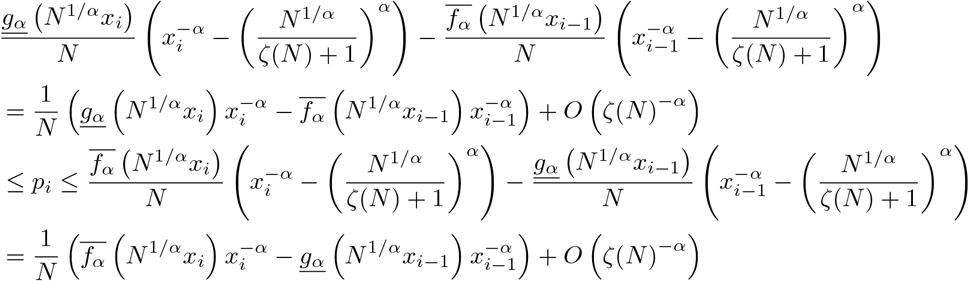

Moreover, *p* ≡ 1 − *p*_1_ −… − *p*_*j*_ = 1 − P (*Y*_1_ ≥ *x*_*j*_|*E*). For positive integers *n*_1_,*…, n*_*j*_ we then have

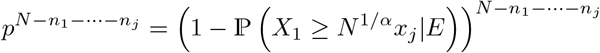

By (27) it then holds that

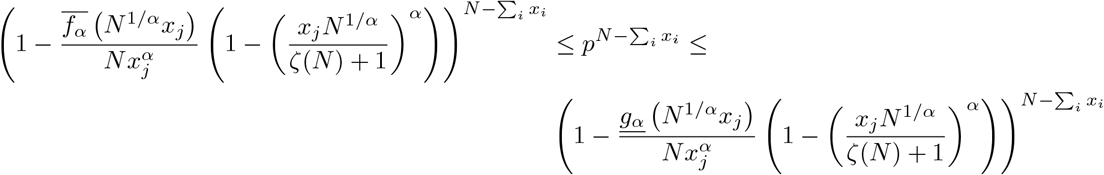

Hence, recalling that *ζ*(*N* )*/N* ^1*/α*^ → ∞ by assumption,

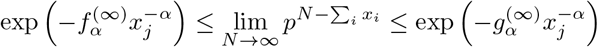

It follows that

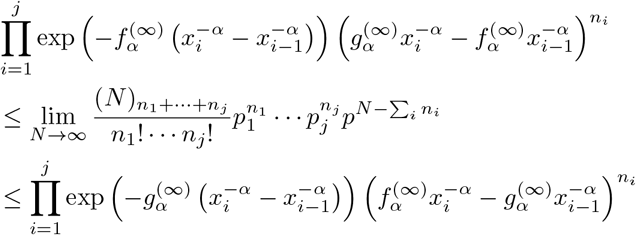

Taking 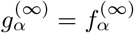 we then have that 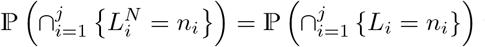 where 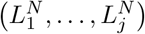 and (*L*_1_,…, *L*_*j*_) are as in the proof of [Schweinsberg, 2003, Lemma 20]. The Poisson-Dirichlet(*α*, 0)-distribution can be constructed from the ranked points *Z*_1_ ≥ *Z*_2_ ≥ … of a Poisson point process on (0, ∞) with characteristic measure 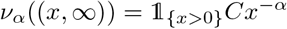 where the law of 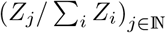 is the Poisson-Dirichlet distribution with parameter (*α*, 0). Let *t:* > 0 and choose *δ* > 0 as in the proof of [Schweinsberg, 2003, Lemma 21] so that *δ* → 0 as *ϵ* → 0. We can use the upper bound in (44) to show that

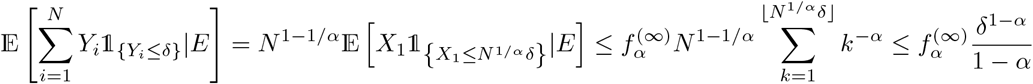

By the arguments in the proof of [Schweinsberg, 2003, Lemma 21] we see that 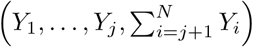 converges weakly to 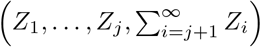 as *N* → ∞. Write 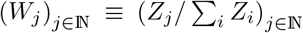 Suppose 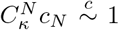, and that 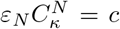 for some *c* > 0 fixed, and for each *N* ∈ ℕ. Then for all *k*,…,*k* ≥ 2, *r* ∈ ℕ, and with Ξ_*α*_ as in Definition 2.6 (recall *C*_*κ*_ from (29b)),

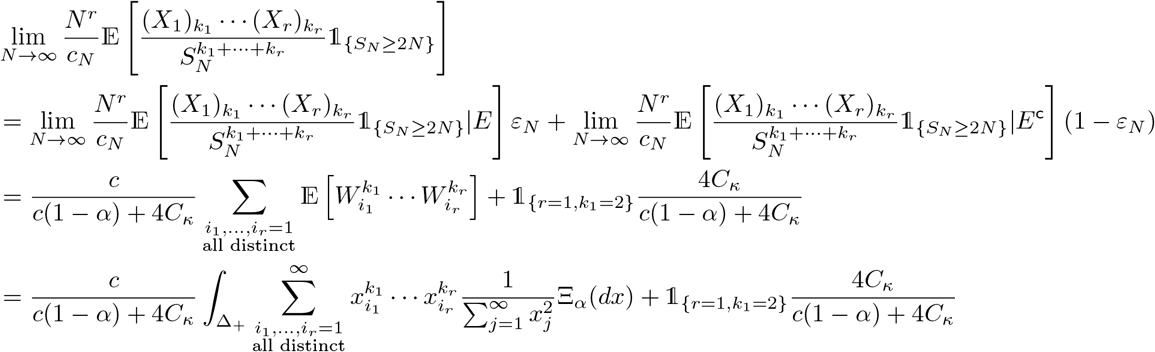

It remains to verify that 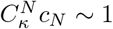. The same arguments as for [Schweinsberg, 2003, Equation 77] give that

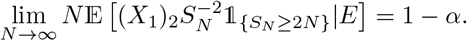

We also have that Given that 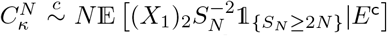; it follows that 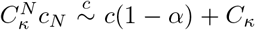 as *N* → ∞. Given that 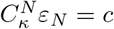, it follows from [Schweinsberg, 2003, Lemma 6, Equation 16] that

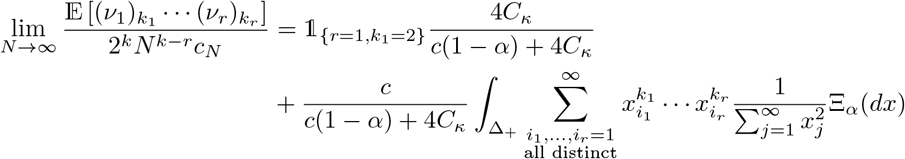

To get the full rate of a particular combination of up to 4*r* simultaneous mergers from *k*_1_,…, *k*_*r*_ group sizes it remains to sum over all the ways in which such mergers can occur. We omit this here; an example when *r* = 1 can be seen in connection with the Ω-*δ*_0_-Beta(*γ*, 2 − *α, α*)-coalescent. Hence, the limits in (17) exist for all *r* ∈ N and *k*_1_,…, *k*_*r*_ ≥ 2, and the proof of Case 3 of Theorem A is complete. □

### 6.2 Proof of Theorem B

In this section we give a proof of Theorem B.

First, we identify conditions on 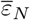 for *m*_∞_ < ∞ (recall (21)) to hold.

#### Lemma 6.14

(Finite *m*_∞_). *Suppose the conditions of Theorem B are in force, and that*

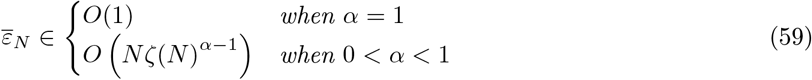

*It then holds that* lim sup_*N* →∞_ *m*_*N*_ < ∞.

*Proof of Lemma 6.14*. Recall the events *E*_1_ and *E*^c^ from Definition 3.10. When 0 < *α* < 1 we see, using the upper bound in (44) and (45),

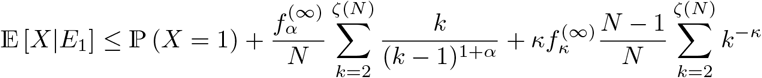

Since ∑_*k*≥1_ *k*^−*κ*^ converges for *κ* ≥ 2 it suffices to check that

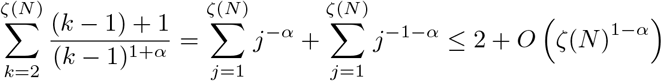

With *X* denoting the random number of potential offspring of an arbitrary parent pair, the result for 0 < *α* < 1 now follows from the relation 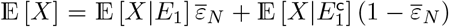. The result for *α* = 1 is obtained in the same way (here we omit the details).

We now check that *S*_*N*_ */*(*Nm*_*N*_ ) → 1 almost surely as *N* → ∞ (recall Lemma 6.8).

#### Lemma 6.15

(*S*_*N*_ */*(*Nm*_*N*_ ) → 1 almost surely). *Under the conditions of Theorem B, with* 1 < *r* < 2 *fixed, and ε*_*N*_ *fulfilling* (59) *in Lemma 6.14 where*

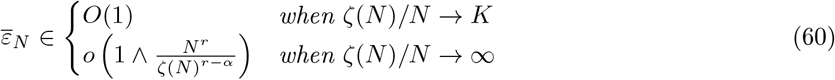

*It then holds that S*_*N*_ */*(*Nm*_*N*_ ) → 1 *almost surely as N* → ∞.

Then

*Proof of Lemma 60*. We will use the approach of [Panov, 2017, Proof of Theorem 2.2i]. Choosing 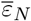 as in (60) ensures that lim sup *N*_→∞_ *m*_*N*_ < ∞ by Lemma 6.14. Define, for 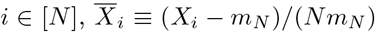. Then

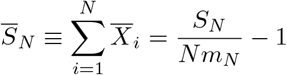

Suppose 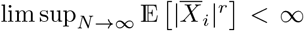 for some 1 < *r* < 2. It then holds by [von Bahr and Esseen, 1965, Theorem 2]

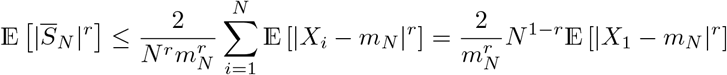

We will now show that 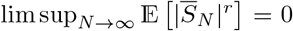. The upper bound in (44) and (45), and the inequality |*x* + *y*| ^*s*^ *≤*2^*s*−1^ (| *x*| ^*s*^ + |*y*| ^*s*^) for any *x, y* ℝ ∈and *s≥* 1 [Athreya and Lahiri, 2006, Proposition 3.1.10*(iii)*, Equation 1.12] combine to give

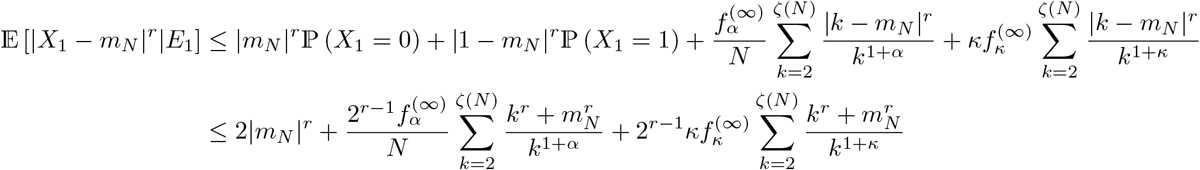

Since 0 < *α* ≤ 1, 1 < *r* < 2, and 2 ≤ *κ*, we see

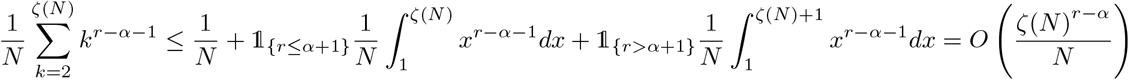

Then E [|*X*_1_ − *m*_*N*_ |^*r*^ |*E*_1_] = *O (*1+ *N* ^−1^*ζ*(*N* )^*r*−*α*^) so that with 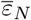 as in (60) it holds that, for 1 < *r* < 2,

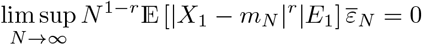

One checks in the same way that 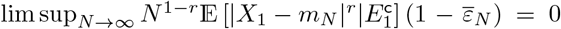(here we omit the details). We have shown that there is an *r* > 1 such that 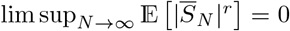. For *i* > *N* take 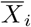 to be independent with mean 0. It follows that 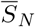 converges in probability to 0. The 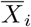 are independent for all *i*∈ n, and so 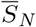 converges almost surely to 0 (e.g. [Athreya and Lahiri, 2006, Theorem 8.3.3]). Hence, 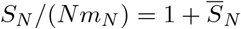 converges almost surely to 1.

We now verify (31) (i.e. 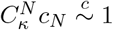 where 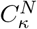 is as in (23)).

#### Lemma 6.16

(Verifying (31)). *Equation* (31) *is in force under the conditions of Theorem B when taking*

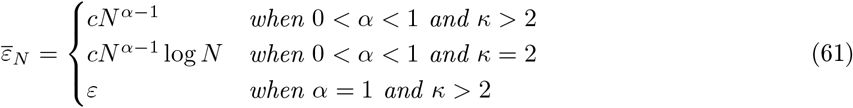

*where* 0 < *ε* < 1 *and c* > 0 *are fixed*.

*Proof of Lemma 6.16*. The proof follows by similar arguments as the proof of Lemma 6.10. Given *ϵ:* > 0 fixed we use Lemma 60 to get (55) with *Y*_−_ and *Y*_+_ as in (54). Then, by Definition 3.10,

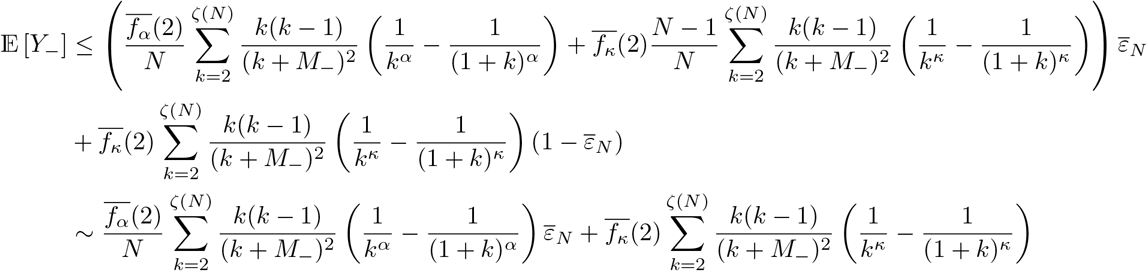

The lower bound on E [*Y*_+_] is of the same form as the upper bound on E[*Y*_−_] except with <u>*g*_*a*_</u>(2) replacing 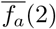. The lemma then follows from Propositions 6.4 and 6.5 as in the proof of Lemma 6.10, and after checking that (56) holds (here we omit the straightforward calculations).

The following lemma checks that a necessary and sufficient condition for convergence to the Kingman coalescent holds under the conditions of Case 1 of Theorem B; the proof follows by similar arguments as the proof of Lemma 6.11, and is omitted.

#### Lemma 6.17

(Convergence to the Kingman coalescent). *Under the conditions of Case 1 of Theorem B and with* 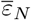 *as in* (61) *it holds that* 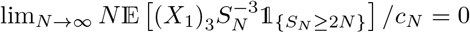

Under the conditions of Case 2 of Theorem B simultaneous mergers will not be generated by the occurrence of two or more large families in any given generation in a large population (as *N* → ∞ ). The proof follows similar arguments as the proof of Lemma 6.12 and is omitted.

#### Lemma 6.18

(The probability of two or more large families vanishes as *N* → ∞). *Under the conditions of Case 2 of Theorem B and with* 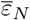 *as in* (61) *it holds that*

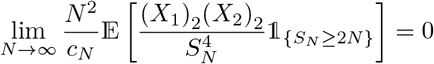

*Proof of Theorem B*. Case 1 of Theorem B is Lemma 6.17 and Lemma 6.2. Convergence to the Kingman coalescent follows as in the proof of Lemma 6.11.

Case 2 of Theorem B follows by identical arguments as for Case 2 of Theorem A. The proof of Theorem B is complete.

## A Computing 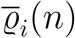 for the Ω-Beta(2 − *β, β*)-Poisson-Dirichlet(*α*, 0)-coalescent

In this section we briefly present an algorithm for computing the empirical mean 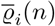 of annealed relative branch lengths *R*_*i*_(*n*) (recall (34), (35a), and (35b)) of gene genealogies when governed by the Ω-Beta(2 −*β, β*)-Poisson-Dirichlet(*α*, 0)-coalescent.

Fix 1 < *β* < 2, and 1*/β* < *α* < 1. The Beta(2 − *β, β*)-Poisson-Dirichlet(*α*, 0)-coalescent on the partitions of [*n*] is a continuous-time Ξ-coalescent with measure Ξ of the form (recall Δ_+_ from (9) and Ξ_*α*_ from Definition 2.6)

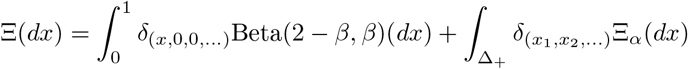

The transition rates are

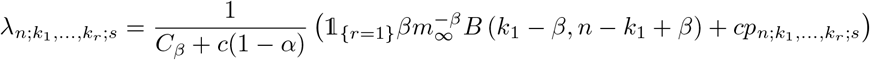

where 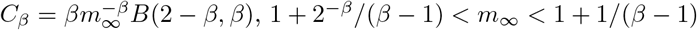, and *p*_*n*;*k*1,…,*kr* ;*s*_ is as in (13).

Consider a diploid panmictic population evolving according to Definition 2.3 and Definition 3.4. The calculations leading to Cases 2 and 3 of Theorem A taking *ζ*(*N* )*/N* ^1*/α*^ → ∞ and 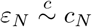 where 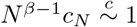 all as *N* → ∞ then lead to the continuous-time Ω-Beta(2 − *β, β*)-Poisson-Dirichlet(*α*, 0)-coalescent.

### Definition A.1

(The Ω-Beta(2 − *β, β*)-Poisson-Dirichlet(*α*, 0)-coalescent). *The* Ω*-Beta*(2 −*β, β*)*-Poisson-Dirichlet*(*α*, 0)*-coalescent, for* 1 < *β* < 2 *and* 1*/β* < *α* < 1, *is the Beta*(2 −*β, β*)*-Poisson-Dirichlet*(*α*, 0)*-coalescent where the blocks in each group split among four subgroups independently and uniformly at random, and the blocks assigned to the same subgroup are merged*.

### Remark A.2

(The range of *α* in Definition A.1). *The range of α in Definition A.1 guarantees that the mean m*_∞_ < ∞, *as one can easily check using the upper bound in* (44) *in Lemma 6.1 (Recall also Remark 3.5)*.

The measure driving the Ω-Beta(2 − *β, β*)-Poisson-Dirichlet(*α*, 0)-coalescent is of the form

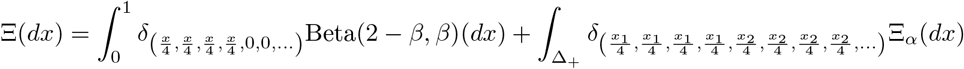

Figure A1 contains examples of 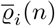 (35a) predicted by the Ω-Beta(2 − *β, β*)-Poisson-Dirichlet(*α*, 0)-coalescent. The graphs in Figure A1c indicate that the Ω-Beta(2 − *β, β*)-Poisson-Dirichlet(*α*, 0)-coalescent can predict a non-monotone (U-shaped) site-frequency spectrum with peaks around the middle of the spectrum. However, Figures A1a and A1b indicate there are cases where the site-frequency spectrum is U-shaped without the peaks, and resembling the site-frequency spectrum observed in population genomic data from the diploid Atlantic cod [Árnason et al., 2023].

## B Computing 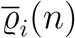 for the Ω-*δ*_0_-Beta(*γ*, 2 − *α, α*)-coalescent

In this section we briefly describe the algorithm for sampling from the Ω-*δ*_0_-Beta(*γ*, 2 − *α, α*) coalescent and computing the empirical mean 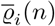 of *R*_*i*_(*n*) (recall (34), (35a), and (35b)). Figure 2 records examples of 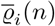.

Let *λ*_*m*_ := Σ _*m*≥*k*1≥…≥*kr* ≥2;_ Σ_*j kj* ≤*m*_ *λ*_*m*;*k*1,…,*kr* ;*s*_ denote the total jump rate out of *m* blocks and *p*_*m*_(*k*) = *λ*_*m*;*k*1,…,*kr* ;*s*_*/λ*_*m*_ the probability of seeing (ordered) merger sizes *k* = (*k*_1_,…, *k*_*r*_) where 2 ≤ *k*_1_ +… + *k*_*r*_ ≤ *m*.

Let

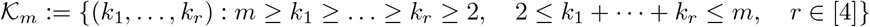

denote the set of all possible mergers when there are *m* blocks and *j*_*m*_ : *K*_*m*_ → [#*K*_*m*_] a map assigning unique indexes to the mergers. Let *F*_*m*_ : [#*K*_*m*_] → [0, 1] denote the cumulative probability mass function for the merger sizes where 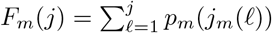.

Let (*l*_*i*_(*n*),…,*l* _*n*−1_(*n*)) denote the realised branch lengths and (*b*_1_,…, *b*_*m*_) the current block sizes where *b*_*j*_ ∈ [2*n*] and Σ_*j*_ *b*_*j*_ = 2*n*.

1. (r_1_(*n*),…, r_2*n*−1_(*n*)) ← (0,…, 0)
2. for each of *M* experiments
  a. (*ℓ*_1_(*n*),…, *ℓ*_2*n*−1_(*n*)) ← (0,…, 0)
  b. set the current number of blocks *m* ← 2*n*
  c. (*b*_1_,…, *b*_*m*_) ← (1,…, 1)
  d. **while** *m* > 1:
    i. sample a random exponential *t* with rate *λ*_*m*_
    ii. ℓ_*b*_(*n*) ← *t* + *ℓ* _*b*_(*n*) for *b* = *b*_1_,…, *b*_*m*_
    iii. sample merger sizes inf {*j* ∈ [|*K*_*m*_|] : *U* ≤ *F*_*m*_(*j*)} where *U* a standard random uniform
    iv. merge blocks according to sampled merger sizes *k*_1_,…, *k*_*r*_
    v. *m* ← *m* − *k*_1_ −· · · − *k*_*r*_ + *r*
  e. r_*i*_(*n*) ← r_*i*_(*n*)+*l* _*i*_(*n*)*/* Σ_*j*_ *l*_*j*_(*n*) for *i* ∈ [2*n* − 1]
3. return an approximation 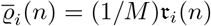 of E for [*R*_*i*_(*n*)] for *i* = 1, 2,…, 2*n* − 1

**Figure A1:**
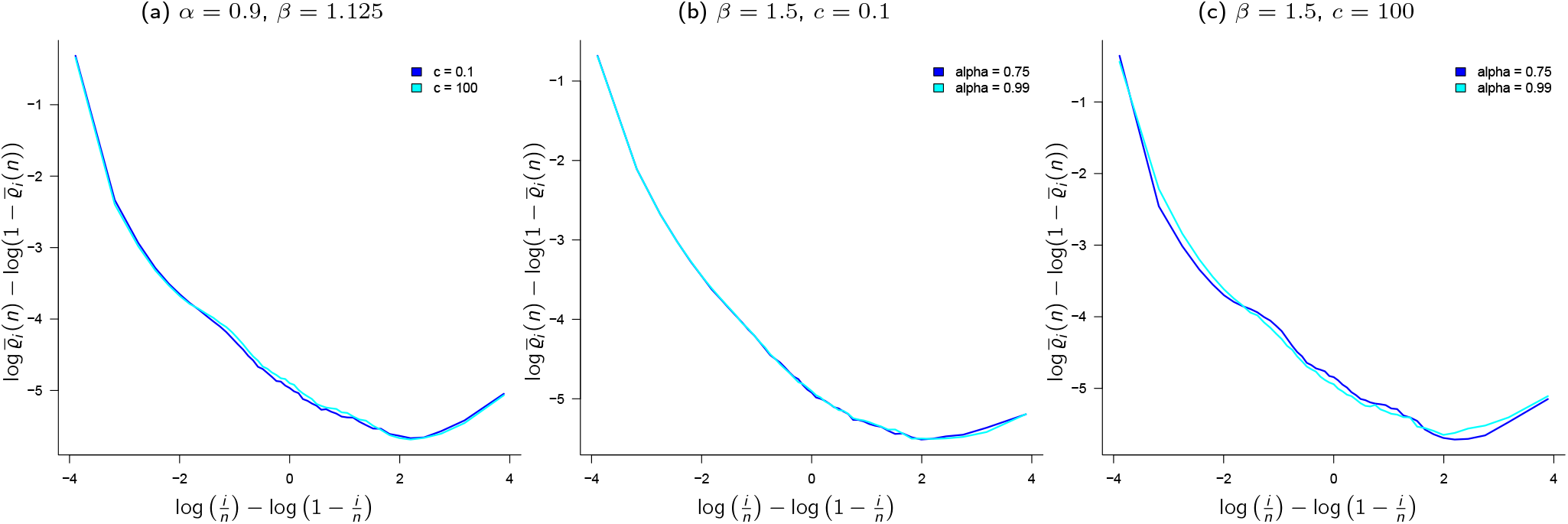
The Ω-Beta(2 − *β, β*)-Poisson-Dirichlet(*α*, 0)-coalescent. Examples of 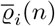 (recall (34), (35a) and (35b)) when {*ξ*^*n*^} is the Ω-Beta(2 − *β, β*)-Poiss on-D irichlet(*α*, 0)-coalesc ent as in Definition A.1 for *n* = 50, *c, α, β* as shown, and approximating *m*_∞_ with (2+ 1+ 2^1−*β*^) */*(*β* − 1) */*2. The scale of the ordinate (y-axis) may vary between graphs; results from 10^5^ experiments

## C Computing 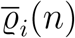 for the Ω-*δ*_0_-Poisson-Dirichlet(*α*, 0)-coalescent

In this section we briefly describe the algorithm for computing the empirical mean 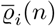 of *R*_*i*_(*n*)(recall (34), (35a), and (35b)) as predicted by the Ω-*δ*_0_-Poisson-Dirichlet(*α*, 0) coalescent (recall Definition 2.9). In Figure C1 we record examples of 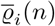 for sample size *n* = 50.

1. (r_1_(*n*),…, r_2*n*−1_(*n*)) ← (0,…, 0)
2. for each of *M* experiments:
  a. (*ℓ*_1_(*n*),…,*ℓ* _2*n*−1_(*n*)) ← (0,…, 0)
  b. *m* ← 2*n*
  c. (*b*_1_,…, *b*_2*n*_) ← (1,…, 1)
  d. **while** *m* > 1:
    i. *t* ← 0
    ii. until a merger occurs (at least one subgroup with at least 2 blocks):
      A. sample group sizes *k*_1_,…, *k*_*r*_ with rate (30)
      B. assign the blocks in each group to one of 4 subgroups independently and uniformly at random
      C. *t* ← *t* + Exp(*λ*_*m*_) where *λ*_*m*_ is the sum of the group sizes rates (30) for *m* blocks
    iii. ℓ_*b*_(*n*) ← *t* + *l*_*b*_(*n*) for *b* = *b*_1_,…, *b*_*m*_
    iv. merge blocks in the same subgroup and update *m*
  e. update r_*i*_(*n*) given one realisation ( *x2113*_1_(*n*),…, *x2113*_2*n*−1_(*n*)) of branch lengths:

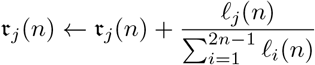
3. return an approximation 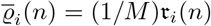 for of E [*R*_*i*_(*n*)] for *i* = 1, 2,…, 2*n* − 1

**Figure C1:**
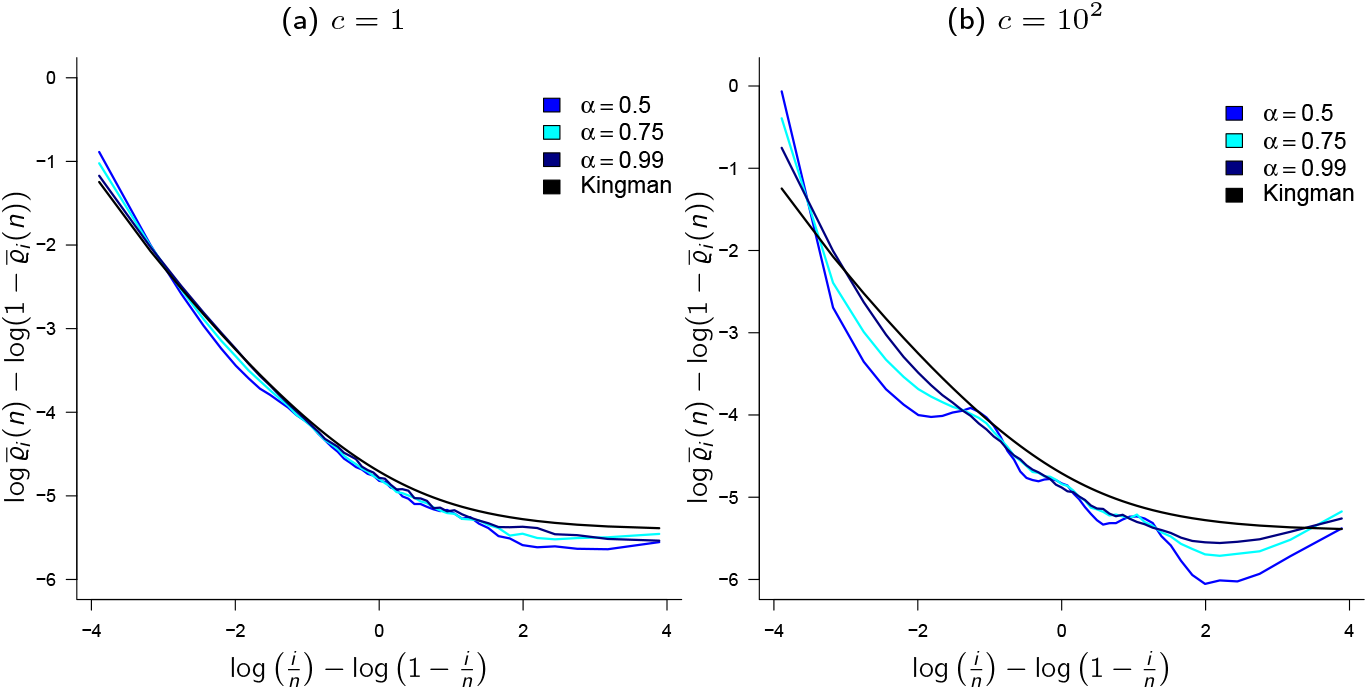
The Ω-*δ*_0_-Poisson-Dirichlet(*α*, 0) coalescent. Empirical mean 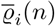 of *R*_*i*_(*n*) (recall (34), (35a)) when{*ξ*^*n*^}is the Ω-*δ*_0_-Poisson-Dirichlet(*α*, 0) coalescent (Definition 2.9); for sample size *n* = 50, *κ* = 2, *c* and *α* as shown; results from 10^5^ experiments. The scale of the ordinate (y-axis) may vary between graphs

## D Computing 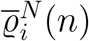

In this section we briefly describe the algorithm for computing 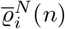 (35b), the empirical mean of 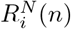 (recall (34)) from a finite number of annealed gene genealogies. Suppose our sample comes from a population evolving according to Def 2.3. Since we are only estimating branch lengths it suffices to keep track of the current block sizes. However, our population is diploid and so we need to keep track of the pairing of blocks in diploid individuals. Thus, we record the current sample configuration in the form of pairs of block sizes. At time 0 we have *n* pairs of {1, 1}, {{1, 1 },…, {1, 1}} ; a common ancestor is reached upon entering the configuration {{2*n*, 0}} . Thus, the sample configuration can be seen as a record of the current marked individuals, where a ‘marked’ individual carries at least one ancestral block. Single-marked individuals carrying one marked block are recorded as {*b*, 0}. Let *n* denote the number of sampled diploid individuals so that 2*n* is the number of sampled gene copies.

1. 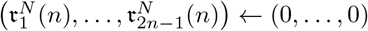 realised relative branch lengths
2. for each of *M* experiments
  a. 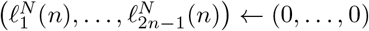
  b. initialise block sizes {{*b*_1_, *b*_2_},…, {*b*_2*n*−1_, *b*_2*n*_}} ← {{1, 1},…, {1, 1}}
  c. **while** all block sizes are smaller than 2*n*:
    i. 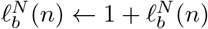 for current block sizes *b* = *b*_1_,…, *b*_*m*_
    ii. sample numbers of potential offspring *X*_1_,…, *X*_*N*_
    iii. given *X*_1_,…, *X*_*N*_ assign marked diploid individuals to families
    iv. merge blocks and record the new block sizes and configuration (pairing in individuals)
  d. given branch lengths 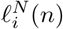 update 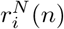 for *i* = 1, 2,…, 2*n* − 1

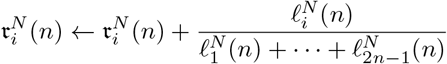
3. return an approximation 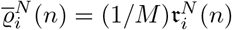 of 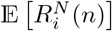

## E Computing 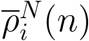

In this section we briefly describe the algorithm for computing 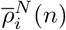, the empirical mean of quenched relative branch lengths 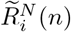 (see also § 4.2). Let A^(*N,n*)^ = (*A*_*i*_(*g*)) _*i*∈[4*N*],*g*∈ ℕ∪{0}_ denote a realised ancestry, recording the ancestral relations of the 4*N* gene copies (chromosomes) in the population. In each generation there are 4*N* gene copies, and each chromosome occupies a level. If the chromosome on level *l* at time *g* produces *k* surviving copies then 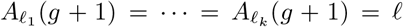, and if *A*_*i*_(*g*) = *A*_*j*_(*g*) then the chromosomes on level *i* resp. *j* at time *g* share an immediate ancestor. In this way the ancestry records the ancestral relations of 4*N* chromosomes, where *N* is the number of parent pairs producing potential offspring in each generation.

The algorithm here is different from the one for approximating 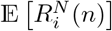 described in Appendix D in that here we generate gene genealogies by recording ancestral relations. In each experiment the population evolves forward in time, the ancestral relations are recorded and stored in A^(*N,n*)^, and every now and then we randomly sample *n* diploid individuals (2*n* gene copies); this process is repeated until the sampled gene copies are found to have common ancestor; the gene tree of the sampled gene copies is then fixed and the branch lengths are read off the fixed tree (a sample whose gene copies are without a common ancestor and so with an incomplete gene tree is discarded). This process is repeated a given number of times, each time starting from scratch with a new population. Averaging over population ancestries gives 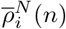 (see Figures 4 and E1 for examples).

1. 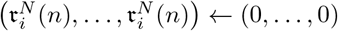
2. for each of *M* experiments:
  a. initialise the ancestry *Al* (0) = *l* for *l*= 1, 2,…, 4*N*
  b. 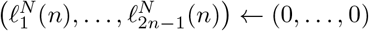
  c. **until** a complete sample tree is found:
    a. add to A^(*N,n*)^:
      I. sample numbers of potential offspring *X*_1_,…, *X*_*N*_
      II. assign chromosomes to each potential offspring; the chromosomes of family *i* will occupy levels 4*i*, 4*i* + 1, 4*i* + 2, 4*i* +3
      III. sample 2*N* surviving offspring
    b. sample uniformly at random and without replacement 2*n* levels
    c. check if the sample tree is complete
  d. read the branch lengths 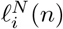 off the fixed complete tree
  e. update 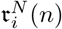 for *i* = 1, 2,…, 2*n* − 1

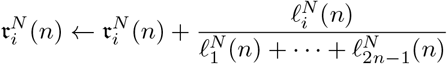
3. return an approximation 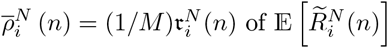

Denote the *M* independently generated ancestries by 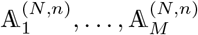 and write 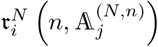 for relative branch lengths read off a complete tree as recorded in 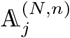. Then for *i* = 1, 2,…, 2*n* − 1

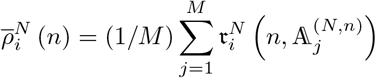

In Figures E1 and 4 are examples comparing 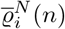 (see § D) and 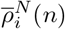; the graphs show the same pattern.

**Figure E1:**
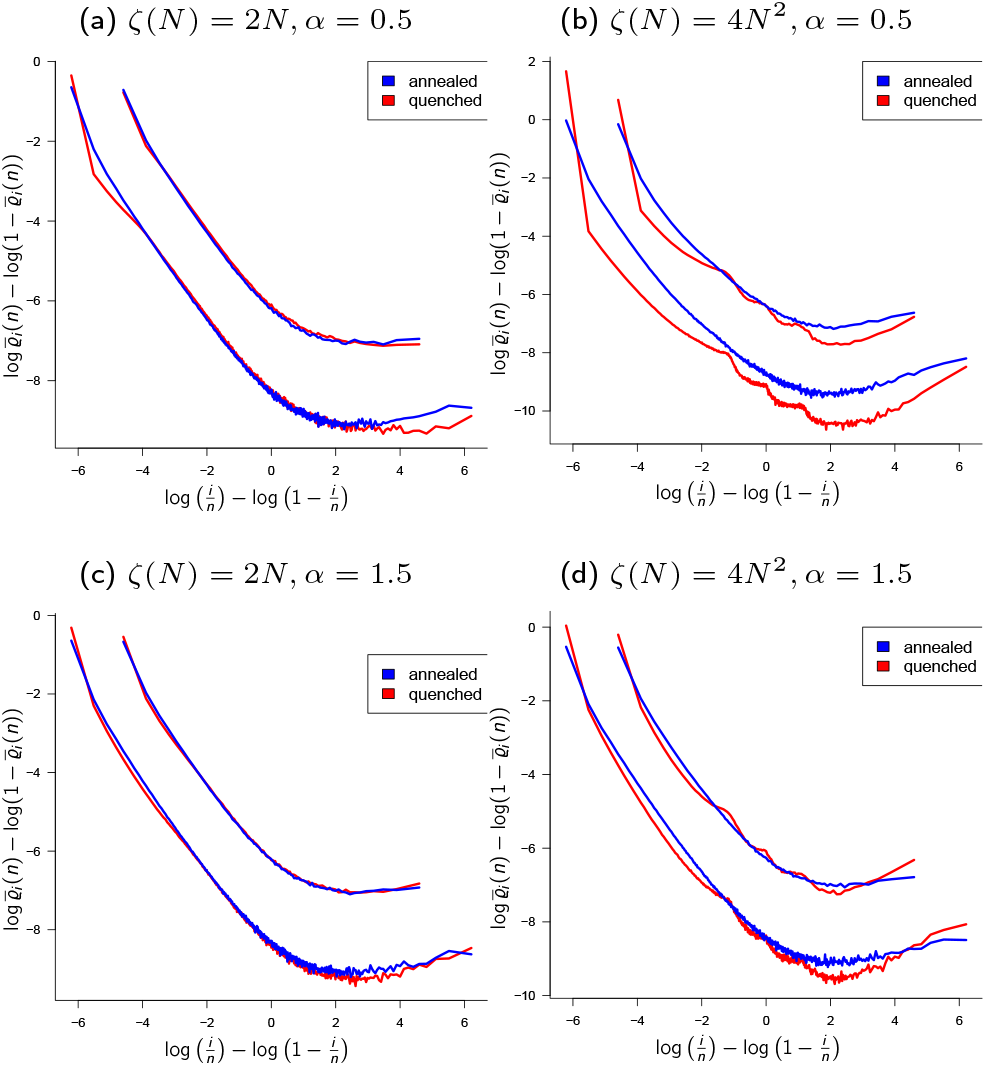
Comparing 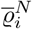 and 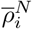. Annealed 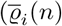, the empirical mean of 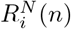, recall (34), (35a) and (35b), blue lines) and quenched 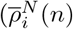, the empirical mean of 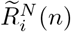, red lines) means when the population evolves according to Definitions 2.3 and 3.4, and (36), for *N* = 125 (population size 2*N* of diploid individuals), *α* and upper bound *ζ*(*N* ) as shown, *ε*_*N*_ = 0.1, *κ* = 2; sample size *n* = 50 and *n* = 250 diploid individuals; results from 10^5^ experiments

## Declarations

### Ethical Statement

The author has no conflict of interest to declare that are relevant to the content of this article

### Data availability

The code (C/C++) to generate the numerical results is freely available at https://github.com/eldonb/gene_genealogies_diploid_pops_sweepstakes

